# Insights from a general, full-likelihood Bayesian approach to inferring shared evolutionary events from genomic data: Inferring shared demographic events is challenging

**DOI:** 10.1101/679878

**Authors:** Jamie R. Oaks, Nadia L’Bahy, Kerry A. Cobb

**Affiliations:** Department of Biological Sciences & Museum of Natural History, Auburn University, Auburn, Alabama 36849; Department of Biology, University of Massachusetts, Amherst, Massachusetts 01003

**Keywords:** phylogeography, biogeography, Bayesian model choice, Dirichlet-process prior

## Abstract

Factors that influence the distribution, abundance, and diversification of species can simultaneously affect multiple evolutionary lineages within or across communities. These include changes to the environment or inter-specific ecological interactions that cause ranges of multiple species to contract, expand, or fragment. Such processes predict temporally clustered evolutionary events across species, such as synchronous population divergences and/or changes in population size. There have been a number of methods developed to infer shared divergences or changes in population size, but not both, and the latter has been limited to approximate methods. We introduce a full-likelihood Bayesian method that uses genomic data to estimate temporal clustering of an arbitrary mix of population divergences and population-size changes across taxa. Using simulated data, we find that estimating the timing and sharing of demographic changes tends to be inaccurate and sensitive to prior assumptions, which is in contrast to accurate, precise, and robust estimates of shared divergence times. We also show previous estimates of co-expansion among five Alaskan populations of threespine sticklebacks (*Gasterosteus aculeatus*) were likely driven by prior assumptions and ignoring invariant characters. We conclude by discussing potential avenues to improve the estimation of synchronous demographic changes across populations.

## 1 Introduction

A primary goal of ecology and evolutionary biology is to understand the processes influencing the distribution, abundance, and diversification of species. Many biotic and abiotic factors that shape the distribution of biodiversity across a landscape are expected to affect multiple species. Abiotic mechanisms include changes to the environment that can cause codistributed species to contract or expand their ranges and/or become fragmented (Hairston et al., 1960; Wegener, 1966; Avise et al., 1987; Knowles and Maddison, 2002). Biotic factors include inter-specific ecological interactions such as the population expansion of a species causing the expansion of its symbionts and the population contraction and/or fragmentation of its competitors (Lotka, 1920; Volterra, 1926; Hairston et al., 1960; Hardin, 1960; Begon et al., 1996; Lunau, 2004). Such processes predict that evolutionary events, such as population divergences or demographic changes, will be temporally clustered across multiple species. As a result, statistical methods that infer such patterns from genetic data allow ecologists and evolutionary biologists to test hypotheses about such processes operating at or above the scale of communities of species.

Recently, researchers have developed methods to infer patterns of temporally clustered (or “shared”) evolutionary events, including shared divergence times among pairs of populations (Hickerson et al., 2006, 2007; Huang et al., 2011; Oaks, 2014, 2019) and shared demographic changes in effective population size across populations (Chan et al., 2014; Xue and Hickerson, 2015; Burbrink et al., 2016; Prates et al., 2016; Xue and Hickerson, 2017; Gehara et al., 2017) from comparative genetic data. To date, no method has allowed the joint inference of both shared divergences and population-size changes. Given the overlap among processes that can potentially cause divergence and demographic changes of populations across multiple species, such a method would be useful for testing hypotheses about community-scale processes that shape biodiversity across landscapes. Here, we introduce a general, full-likelihood Bayesian method that can estimate shared times among an arbitrary mix of population divergences and population size changes (Figure 1).

**Figure 1.**
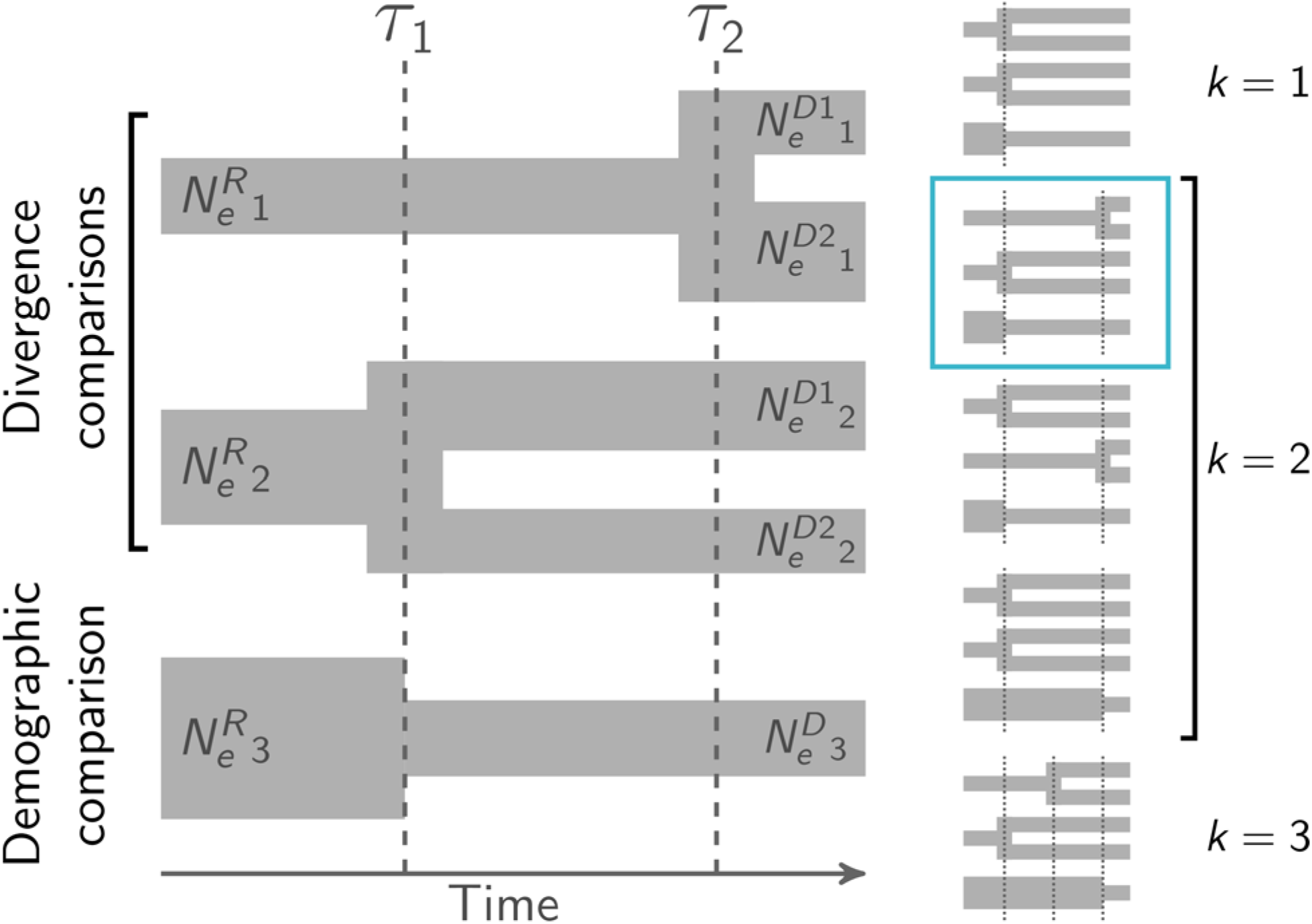
An illustration of the general comparative model implemented in ecoevolity. The top two comparisons are pairs of populations for which we are interested in comparing their time of divergence (“divergence comparisons”). The bottom comparison is a single population for which we are interested in comparing the time of population-size change (“demographic comparison”). With three comparisons, there are five possible event models (i.e., five ways to assign the comparisons to anywhere from one to three event times; Bell, 1934). These five models are illustrated to the right, with the number of events with unique times (*k*) ranging from 1–3; the enlarged example model is indicated with a border. The event time (*τ*_1_ and *τ*_2_) and effective population size 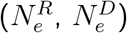 parameters are shown. Event times can be shared among comparisons, but each ancestral and descendant population has a unique effective population size.

Whereas the theory and performance of methods that estimate shared divergence times has been relatively well-investigated (e.g., Oaks et al., 2013; Hickerson et al., 2014; Oaks et al., 2014; Oaks, 2014; Overcast et al., 2017; Oaks, 2019), exploration into the estimation of shared changes in population size has been much more limited. There are theoretical reasons to suspect that estimating shared changes in effective population size is more difficult than divergence times (Myers et al., 2008). The parameter of interest (timing of a demographic change) is informed by differing rates at which sampled copies of a locus “find” their common ancestors (coalesce) going backward in time before and after the change in population size, and this can become unidentifiable in three ways. First, as the magnitude of the change in population size becomes smaller, it becomes more difficult to identify, because the rates of coalescence before and after the change become more similar. Second, as the age of the demographic change increases, fewer of the genetic coalescent events occur prior to the change, resulting in less information about the effective size of the population prior to the change, and thus less information about the magnitude and timing of the populationsize change itself. Third, information also decreases as the age of the demographic change approaches zero, because fewer coalescent events occur after the change.

To explore these potential problems, we take advantage of our full-likelihood method to assess how well we can infer shared demographic changes among populations when using all the information in genomic data. We apply our method to restriction-site-associated DNA sequence (RADseq) data from five populations of three-spine stickleback (*Gasterosteus aculeatus*; Hohenlohe et al., 2010) that were previously estimated to have co-expanded with an approximate Bayesian computation (ABC) approach (Xue and Hickerson, 2015). In stark contrast to shared divergence times, our results show that estimates of shared changes in population size are quite poor across a broad range of simulation conditions. We also find strikingly different estimates of the demographic histories of the stickleback populations depending on whether we include invariant sites in analyses. This alarming result makes sense in light of the inference pathologies exhibited by our analyses of simulated data, where limited information in the data coupled with limited prior knowledge about parameters leads to spurious support for shared demographic changes across populations.

## 2 The model

We extended the model implemented in the software package `ecoevolity` to accommodate two types of temporal comparisons that are specified *a priori* by the investigator:

1. A population that experienced a change from effective population size 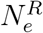 to effective size 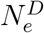 at time *t* in the past. We will refer to this as a *demographic comparison* (Figure 1), and refer to the population before and after the change in population size as “ancestral” and “descendant”, respectively.
2. A population that diverged at time *t* in the past into two descendant populations, each with unique effective population sizes. We will refer to this as a *divergence comparison* (Figure 1).

This allows inference of shared times of divergence and/or demographic change across an arbitrary mix of demographic and divergence comparisons in a full-likelihood, Bayesian framework. Table 1 provides a key to the notation we use throughout this paper.

**Table 1.** A key to some of the notation used in the text.

| Symbol | Description |
| --- | --- |
| $\mathcal{N}$ | The number of comparisons (or taxa); can be an arbitrary mix of populations (comparing timing of demographic change) and/or pairs of populations (comparing timing of divergence). |
| $k$ | The number of events (unique times) across the comparisons. |
| $t_i$ | The time in the past when comparison $i$ either diverged or experienced a change in effective population size. |
| $\tau$ | An event time at which one or more comparisons experienced a divergence or change in effective population size. |
| $\mathcal{T}$ | The event-time model, which comprises the assignment of comparisons to events. |
| $\boldsymbol{\tau}$ | All of the times of the events in the model ($\boldsymbol{\tau} = \tau_1, \dots, \tau_k$). |
| $\alpha$ | The concentration parameter of the Dirichlet process. |
| $n, r$ | The number of copies of a locus sampled from a population, and the number of those copies that are the “red” allele. |
| $\mathbf{n}, \mathbf{r}$ | The allele counts from a comparison (one or two populations). |
| $D_i$ | The allele counts across all characters from comparison $i$. I.e., all of the characters being analyzed for comparison $i$. |
| $m$ | The number of characters collected from a taxon (comparison). |
| $\mathbf{D}$ | All of the data being analyzed, i.e., the character matrices from all comparisons. |
| $g$ | A gene tree with branch lengths. |
| $\mu$ | The rate of mutation. |
| $u$ | Relative rate of mutating from the “red” to “green” state. |
| $v$ | Relative rate of mutating from the “green” to “red” state. |
| $\pi$ | The stationary frequency of the “green” state. |
| $N_e^D$ | The effective size of a descendant population. |
| $N_e^R$ | The effective size of the root (ancestral) population. |
| $R_{N_e^R}$ | The relative effective population size of the root (ancestral) population; relative to the mean of the effective sizes of the descendant populations. |
| $\mathbb{N}_e$ | Shorthand notation for all effective population sizes for a comparison (ancestral and one or two descendant populations). |
| $S$ | The species tree for a comparison. This comprises the effective population sizes and the time of demographic change or divergence. |

### 2.1 The data

As described by Oaks (2019), we assume we have collected orthologous, biallelic genetic characters from taxa we wish to compare. By biallelic, we mean that each character has at most two states, which we refer to as “red” and “green” following Bryant et al. (2012). For each comparison, we either have these data from one or more individuals from a single population, in which case we infer the timing and extent of a population size change, or one or more individuals from two populations or species, in which case we infer the time when they diverged (Figure 1).

For each population and for each character we genotype *n* copies of the locus, *r* of which are copies of the red allele and the remaining *n* − *r* are copies of the green allele. Thus, for each population, and for each character, we have a count of the total sampled gene copies and how many of those are the red allele. Following the notation of Oaks (2019) we will use **n** and **r** to denote allele counts for a character from either one population if we are modeling a population-size change or both populations of a pair if we are modeling a divergence; i.e., **n**, **r** = (*n, r*) or **n**, **r** = {(*n*_1_, *r*_1_), (*n*_2_, *r*_2_)}. For convenience, we will use *D_i_* to denote these allele counts across all the characters from comparison *i*, which can be a single population or a pair of populations. Finally, we use **D** to represent the data across all the taxa for which we wish to compare times of either divergence or population-size change. Note, because the population history of each comparison is modeled separately (Figure 1), characters do not have to be orthologous across comparisons, only within them.

### 2.2 The evolution of characters

We assume each character evolved along a gene tree (*g*) according to a finite-character, continuous-time Markov chain (CTMC) model, and the gene tree of each character is independent of the others, conditional on the population history (i.e., the characters are effectively unlinked). As a character evolves along the gene tree, forward in time, there is a relative rate *u* of mutating from the red state to the green state, and a corresponding relative rate *v* of mutating from green to red (Bryant et al., 2012; Oaks, 2019). The stationary frequency of the green state is then *π* = *u/*(*u* + *v*). We will use *μ* to denote the overall rate of mutation. Evolutionary change is the product of *μ* and time. Thus, if *μ* = 1, time is measured in units of expected substitutions per site. Alternatively, if a mutation rate per site per unit of time is given, then time is in those units (e.g., generations or years).

### 2.3 The evolution of gene trees

We assume the gene tree of each character coalesced within a simple “species” tree with one ancestral root population that, at time *t*, either left one or two descendant branches with different effective population sizes (Figure 1). We will use ℕ_e_ to denote all the effective population sizes of a species tree; 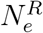 and 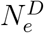 when modeling a population-size change, and 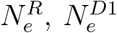, and 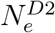 when modeling a divergence. Following Oaks (2019), we use *S* as shorthand for the species tree, which comprises the population sizes and event time of a comparison (ℕ_e_ and *t*).

### 2.4 The likelihood

As in Oaks (2019), we use the work of Bryant et al. (2012) to analytically integrate over all possible gene trees and character substitution histories to compute the likelihood of the species tree directly from a biallelic character pattern under a multi-population coalescent model (Kingman, 1982a,b; Rannala and Yang, 2003); *p*(**n**, **r** | *S*, *μ*, *π*). We only need to make a small modification to accommodate population-size-change models that have a species tree with only one descendant population. Equation 19 of Bryant et al. (2012) shows how to obtain the partial likelihoods at the bottom of an ancestral branch from the partial likelihoods at the top of its two descendant branches. When there is only one descendant branch, this is simplified, and the partial likelihoods at the bottom of the ancestral branch are equal to the partial likelihoods at the top of its sole descendant branch. Other than this small change, the probability of a biallelic character pattern given the species tree, mutation rate, and equilibrium state frequencies (*p*(**n**, **r** | *S*, *μ*, *π*)) is calculated the same as in Bryant et al. (2012) and Oaks (2019).

For a given comparison, we can calculate the probability of all *m* characters for which we have data given the species tree and other parameters by assuming independence among characters (conditional on the species tree) and taking the product over them,

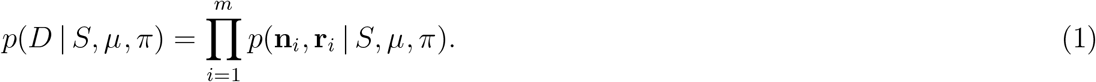

We assume we have sampled biallelic data from 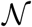 comparisons, which can be an arbitrary mix of (1) two populations or species for which *t* represents the time they diverged, or (2) one population for which *t* represents the time of a change in population size. Assuming independence among comparisons, the likelihood across all 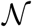 comparisons is simply the product of the likelihood of each comparison,

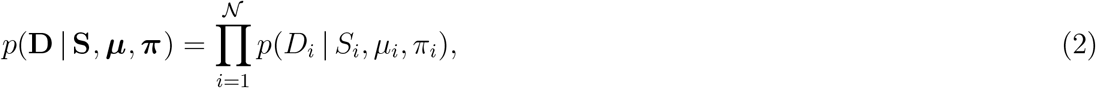

where 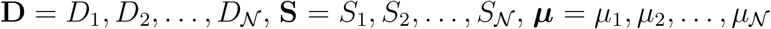 and 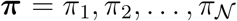. As described in Oaks (2019), if constant characters are not sampled for a comparison, we condition the likelihood for that comparison on only having sampled variable characters.

### 2.5 Bayesian inference

As described by Oaks (2019), to relax the assumption of temporal independence among comparisons, we treat the number of events (population-size changes and/or divergences) and the assignment of comparisons to those events as random variables under a Dirichlet process (Ferguson, 1973; Antoniak, 1974). We use 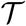 to represent the partitioning of comparisons to events, which we will also refer to as the “event model.” The concentration parameter, *α*, controls how clustered the Dirichlet process is, and determines the probability of all possible 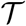 (i.e., all possible set partitions of 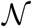 comparisons to 1,2,…,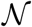 events). We use ***τ*** to represent the unique times of events in 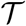. Using this notation, the posterior distribution of our Dirichlet-process model is

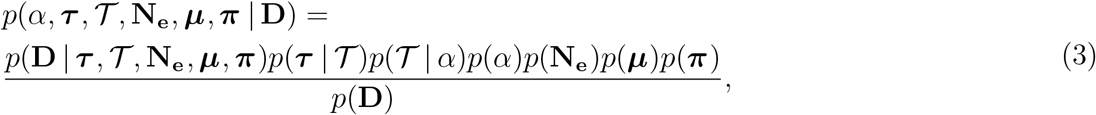

where **N_e_** is the collection of the effective population sizes (ℕ_*e*_) across all of the comparisons.

#### 2.5.1 Priors

##### Prior on the concentration parameter

Our implementation allows for a hierarchical approach to accommodate uncertainty in the concentration parameter of the Dirichlet process by specifying a gamma distribution as a hyperprior on *α* (Escobar and West, 1995; Heath et al., 2011). Alternatively, *α* can also be fixed to a particular value, which is likely sufficient when the number of comparisons is small.

##### Prior on the divergence times

Given the partitioning of comparisons to events, we use a gamma distribution for the prior on the time of each event, 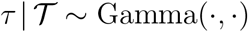.

##### Prior on the effective population sizes

We use a gamma distribution as the prior on the effective size of each descendant population of each comparison. Following Oaks (2019), we use a gamma distribution on the effective size of the ancestral population *relative* to the size of the descendant population(s), which we denote as 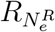. For a comparison with two descendant populations (i.e., a divergence comparison), the prior on the ancestral population size is specified as relative to the mean of the descendant populations. For a comparison with only one descendant population (i.e., a demographic comparison), the prior on the ancestral population is relative to the size of that descendant.

##### Prior on mutation rates

We follow the same approach explained by Oaks (2019) to model mutation rates across comparisons. The decision about how to model mutation rates is extremely important for any comparative phylogeographic approach that models taxa as disconnected species trees (Figure 1; e.g., Hickerson et al., 2006, 2007; Huang et al., 2011; Chan et al., 2014; Oaks, 2014; Xue and Hickerson, 2015; Burbrink et al., 2016; Xue and Hickerson, 2017; Gehara et al., 2017; Oaks, 2019). Time and mutation rate are inextricably linked, and because the comparisons are modeled as separate species trees, the data cannot inform the model about relative or absolute differences in *μ* among the comparisons. We provide flexibility to the investigator to fix or place prior probability distributions on the relative or absolute rate of mutation for each comparison. However, if one chooses to accommodate uncertainty in the mutation rate of one or more comparisons, the priors should be strongly informative. Because of the inextricable link between rate and time, placing a weakly informative prior on a comparison’s mutation rate prevents estimation of the time of its demographic change or divergence, which is the primary goal.

##### Prior on the equilibrium state frequency

Recoding four-state nucleotides to two states requires some arbitrary decisions, and whenever *π* ≠ 0.5, these decisions can affect the likelihood of the model (Oaks, 2019). Because DNA is the dominant character type for genomic data, we assume that *π* = 0.5 in this paper. This makes the CTMC model of character-state substitution a two-state analog of the “JC69” model (Jukes and Cantor, 1969). However, if the genetic markers collected for one or more comparisons are naturally biallelic, the frequencies of the two states can be meaningfully estimated, and our implementation allows for a beta prior on *π* in such cases. This makes the CTMC model of character-state substitution a two-state general time-reversible model (Tavaré, 1986).

#### 2.5.2 Approximating the posterior with MCMC

We use Markov chain Monte Carlo (MCMC) algorithms to sample from the joint posterior in Equation 3. To sample across event models 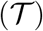 during the MCMC chain, we use the Gibbs sampling algorithm (Algorithm 8) of Neal (2000). We also use univariate and multivariate Metropolis-Hastings algorithms (Metropolis et al., 1953; Hastings, 1970) to update the model, the latter of which are detailed in Oaks (2019).

### 2.6 Software implementation

The `C++` source code for `ecoevolity` is freely available from https://github.com/phyletica/ecoevolity and includes an extensive test suite. From the `C++` source code, two primary command-line tools are compiled: (1) `ecoevolity`, for performing Bayesian inference under the model described above, and (2) `simcoevolity` for simulating data under the model described above. Documentation for how to install and use the software is available at http://phyletica.org/ecoevolity/. We have incorporated help in pre-processing data and post-processing posterior samples collected by `ecoevolity` in the Python package `pycoevolity`, which is available at https://github.com/phyletica/pycoevolity. We used Version 0.3.1 (Commit 9284417) of the `ecoevolity` software package for all of our analyses. A detailed history of this project, including all of the data and scripts needed to produce our results, is available at https://github.com/phyletica/ecoevolity-demog-periments (Oaks et al., 2019a).

## 3 Materials & Methods

### 3.1 Analyses of simulated data

#### 3.1.1 Assessing ability to estimate timing and sharing of demographic changes

We used the `simcoevolity` and `ecoevolity` tools within the `ecoevolity` software package (Oaks, 2019) to simulate and analyze data sets, respectively, under a variety of conditions. Each simulated data set comprised 500,000 unlinked biallelic characters from 10 diploid individuals (20 genomes) sampled per population from three demographic comparisons. We specified the concentration parameter of the Dirichlet process so that the mean number of demographic change events was two (*α* = 1.414216). We assumed the mutation rates of all three populations were equal and 1, such that time and effective population sizes were scaled by the mutation rate. When analyzing each simulated data set, we ran four MCMC chains for 75,000 generations with a sample taken every 50 generations. From preliminary analyses, we calculated the potential scale reduction factor (PSRF; the square root of Equation 1.1 in Brooks and Gelman, 1998) and effective sample size (ESS; Gong and Flegal, 2016) from the four chains for all continuous parameters and the log likelihood using the `pyco-sumchains` tool of `pycoevolity` (Version 0.1.2 Commit 89d90a1). Based on these metrics of MCMC convergence and mixing, we conservatively chose to summarize the last 1000 samples from each chain for a total of 4000 samples of parameter values to approximate the posterior distribution for every simulated data set. When plotting results, we highlight any simulation replicates that have a PSRF > 1.2.

Initially, we simulated data under a variety of settings we thought covered regions of parameter space that are both conducive and challenging for estimating the timing and sharing of demographic changes. However, estimates were quite poor across all our initial simulation conditions (see Supporting Information). In an effort to find conditions under which the timing and sharing of demographic changes could be better estimated, and avoid combinations of parameter values that caused parameter identifiability problems in our initial analyses, we explored simulations under gamma distributions on times and population sizes offset from zero, and with recent demographic event times, (***V***1 – ***V***5, Table 2). When we specify an “offset,” we are right-shifting the entire gamma distribution to have a lower limit of the offset value, rather than zero.

**Table 2.** Simulation and analysis conditions for all simulation-based analyses with three demographic comparisons. The distributions from which parameter values were drawn for simulating data with simcoevolity are given for event times (*τ*), the relative effective size of the root (ancestral) population 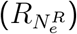, and the effective size of the descendant population 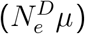, along with the prior distributions used for these parameters when the simulated data sets were analyzed with ecoevolity. When the latter is represented by a dash, this means the prior distribution matched the distribution under which the data were simulated. G(…) and E(…) represent gamma and exponential distributions, respectively, and the parameters of a gamma distribution are given as G^offset^(shape, mean = mean).

| Label | Simulated distribution |  |  | Prior distribution |  |  |
| --- | --- | --- | --- | --- | --- | --- |
| | $\tau$ | $R_{N_e^R}$ | $N_e^D \mu$ | $\tau$ | $R_{N_e^R}$ | $N_e^D \mu$ |
| **Validation simulation conditions** |  |  |  |  |  |  |
| **V1** | $G^{0.0001}(4, \text{mean} = 0.002)$ | $G^{0.05}(5, \text{mean} = 0.25)$ | $G^{0.0001}(4, \text{mean} = 0.0021)$ | - | - | - |
| **V2** | $G^{0.0001}(4, \text{mean} = 0.002)$ | $G^{0.05}(5, \text{mean} = 0.5)$ | $G^{0.0001}(4, \text{mean} = 0.0021)$ | - | - | - |
| **V3** | $G^{0.0001}(4, \text{mean} = 0.002)$ | $G^{0.05}(5, \text{mean} = 4)$ | $G^{0.0001}(4, \text{mean} = 0.0021)$ | - | - | - |
| **V4** | $G^{0.0001}(4, \text{mean} = 0.002)$ | $G^{0.05}(5, \text{mean} = 1)$ | $G^{0.0001}(4, \text{mean} = 0.0021)$ | - | - | - |
| **V5** | $G^{0.0001}(4, \text{mean} = 0.002)$ | $G^{0.05}(50, \text{mean} = 1)$ | $G^{0.0001}(4, \text{mean} = 0.0021)$ | - | - | - |
| **Sensitivity simulation conditions** |  |  |  |  |  |  |
| **S1** | $G^{0.0001}(4, \text{mean} = 0.002)$ | $G^{0.05}(5, \text{mean} = 0.25)$ | $G^{0.0001}(4, \text{mean} = 0.0021)$ | $E(\text{mean} = 0.005)$ | $E(\text{mean} = 2)$ | $G(2, \text{mean} = 0.002)$ |
| **S2** | $G^{0.0001}(4, \text{mean} = 0.002)$ | $G^{3.80}(5, \text{mean} = 4)$ | $G^{0.0001}(4, \text{mean} = 0.0021)$ | $E(\text{mean} = 0.005)$ | $E(\text{mean} = 2)$ | $G(2, \text{mean} = 0.002)$ |
| **S3** | $G^{0.0001}(4, \text{mean} = 0.002)$ | $G^{0.05}(5, \text{mean} = 0.1)$ | $G^{0.0001}(4, \text{mean} = 0.0021)$ | $E(\text{mean} = 0.005)$ | $E(\text{mean} = 2)$ | $G(2, \text{mean} = 0.002)$ |
| **S4** | $G^{0.0001}(4, \text{mean} = 0.002)$ | $G^{9.95}(5, \text{mean} = 10)$ | $G^{0.0001}(4, \text{mean} = 0.0021)$ | $E(\text{mean} = 0.005)$ | $E(\text{mean} = 2)$ | $G(2, \text{mean} = 0.002)$ |

For the mutation-scaled effective size of the descendant population 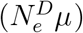, we used an offset gamma distribution with a shape of 4, offset of 0.0001, and mean of 0.0021 after accounting for the offset (Table 2). The mean of this distribution corresponds to an average number of differences per character between individuals in the population (i.e., nucleotide diversity) of 0.0084, which is comparable to estimates from genomic data of populations of zooplankton (Choquet et al., 2019), stickleback fish (Hohenlohe et al., 2010), and humans (Auton et al., 2015). For the distribution of event times, we used a gamma distribution with a shape of 4, offset of 0.0001, and a mean of 0.002 (after accounting for the offset; Table 2). Taken together, this creates a distribution of event times in units of 4*N_e_* generations with a mean of approximately 0.3. For example, for a population of diploids with an effective size of 100,000 individuals and a generation time of one year, the mean is approximately 120,000 years. We chose these distributions to try and balance the number of gene lineages that coalesce after and before the population-size change for the average gene tree. We used the offset values to avoid very small descendant population sizes and very recent times of population-size change, because in our preliminary analyses, both of these conditions caused the timing of events to be essentially nonidentifiable (see Supporting Information).

We chose five different distributions on the relative effective size of the ancestral population (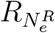; see Table 2 and left column of Figures 2 and 3), which ranged from having a mean 4-fold population-size increase (***V***1) and decrease (***V***3), and a “worst-case” scenario where there was essentially no population-size change in the history of the populations (***V***5). We generated 500 data sets under each of these five conditions (***V***1– ***V***5, Table 2), and analyzed all of them using priors that matched the generating distributions.

**Figure 2.**
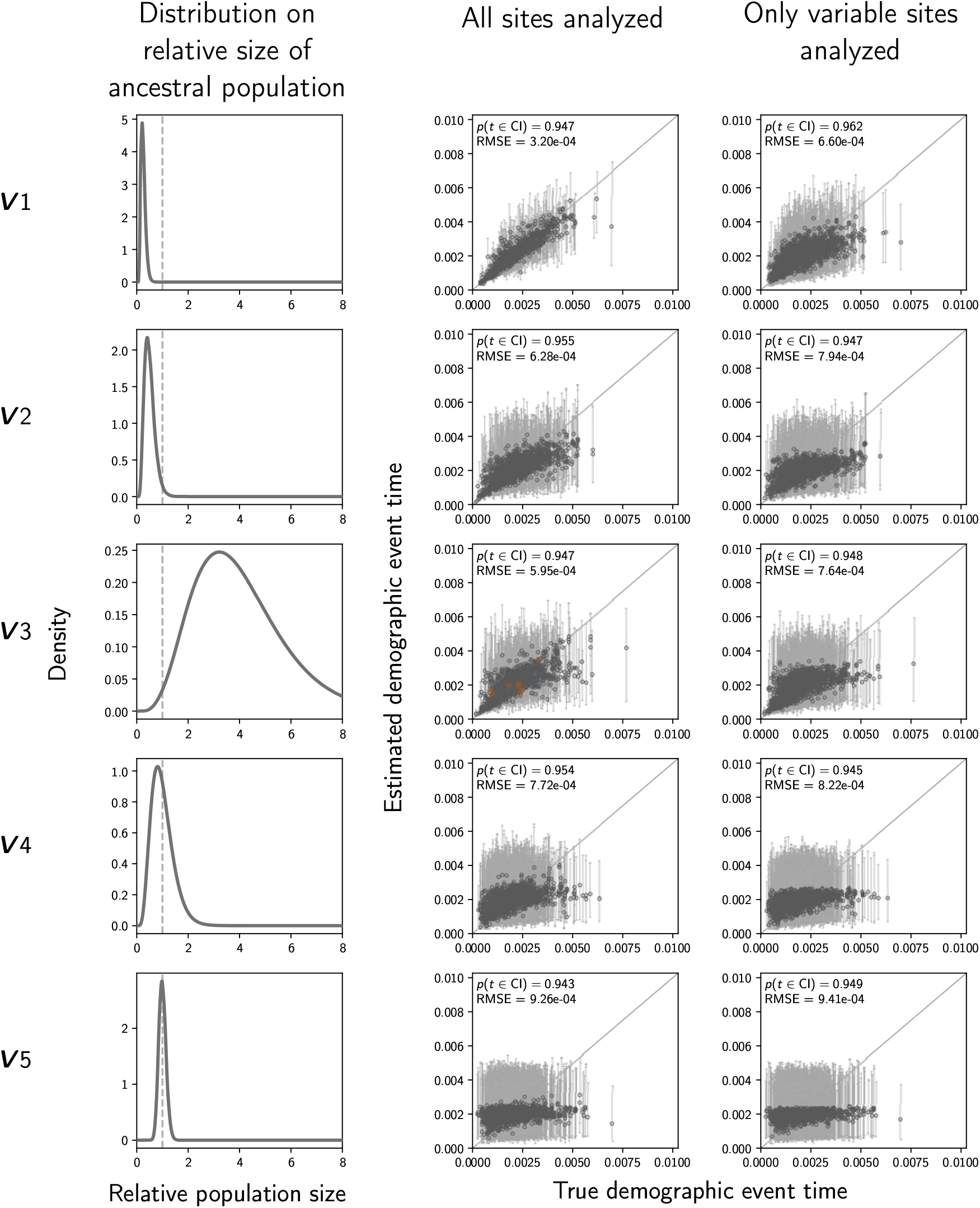
The accuracy and precision of time estimates of demographic changes (in units of expected subsitutions per site) when data were simulated and analyzed under the same distributions (Table 2). The left column of plots shows the gamma distribution from which the relative size of the ancestral population was drawn; this was also used as the prior when each simulated data set was analyzed. The center and right column of plots show true versus estimated values when using all characters (center) or only variable characters (right). Each plotted circle and associated error bars represent the posterior mean and 95% credible interval. Estimates for which the potential-scale reduction factor was greater than 1.2 (Brooks and Gelman, 1998) are highlighted in orange. Each plot consists of 1500 estimates—500 simulated data sets, each with three demographic comparisons. For each plot, the root-mean-square error (RMSE) and the proportion of estimates for which the 95% credible interval contained the true value—*p*(*t* ∈ CI)—is given. We generated the plots using matplotlib Version 2.0.0 (Hunter, 2007).

**Figure 3.**
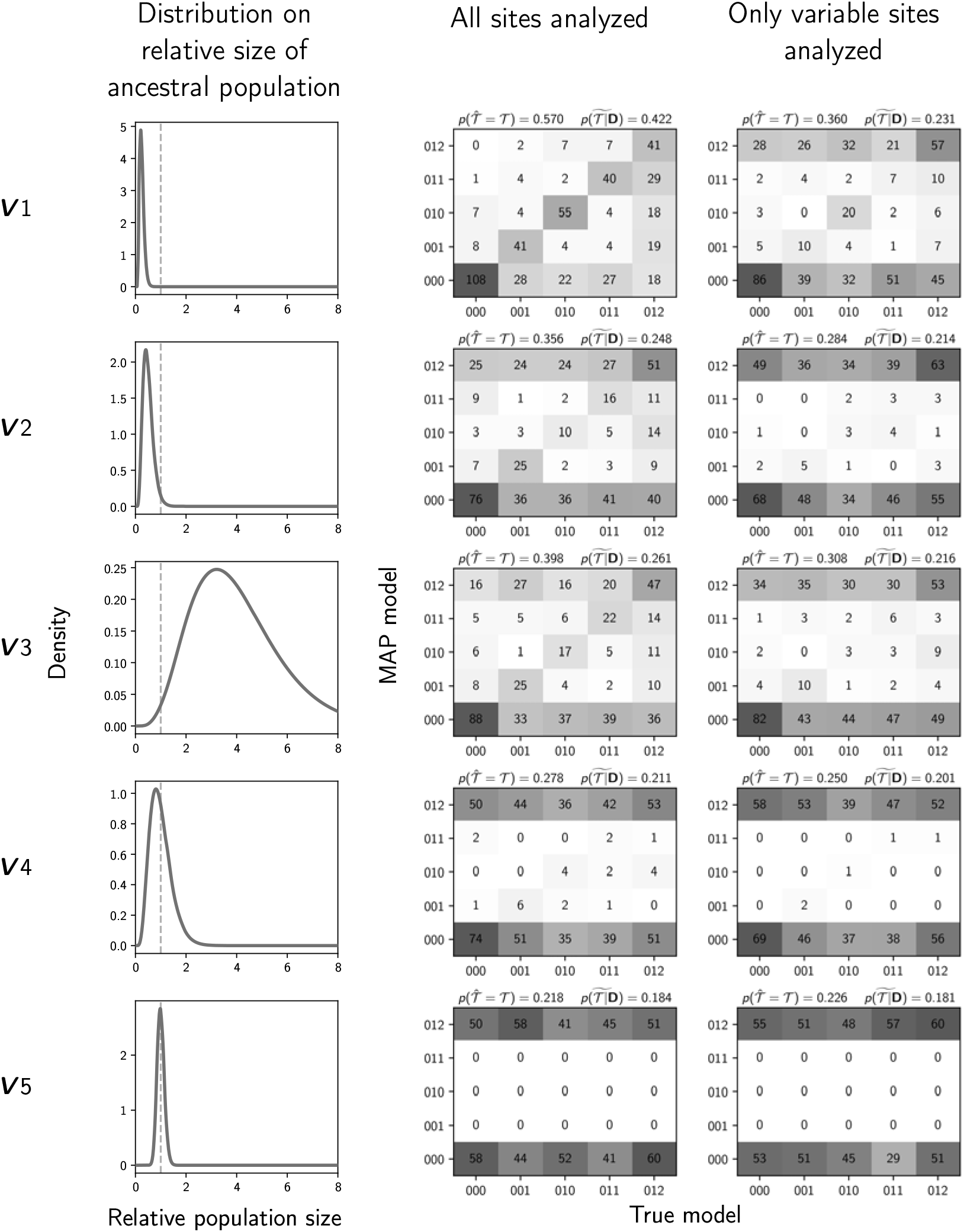
The performance of estimating the model of demographic changes when data were simulated and analyzed under the same distributions (Table 2). The left column of plots shows the gamma distribution from which the relative size of the ancestral population was drawn; this was also used as the prior when each simulated data set was analyzed. The center and right column of plots show true versus estimated models when using all characters (center) or only variable characters (right). Each plot shows the results of the analyses of 500 simulated data sets, each with three demographic comparisons; the number of data sets that fall within each possible cell of true versus estimated model is shown, and cells with more data sets are shaded darker. Each model is represented along the plot axes by three integers that indicate the event category of each comparison (e.g., 011 represents the model in which the second and third comparison share the same event time that is distinct from the first). The estimates are based on the model with the maximum *a posteriori* probability (MAP). For each plot, the proportion of data sets for which the MAP model matched the true model—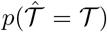—is shown in the upper left corner, and the median posterior probability of the correct model across all data sets—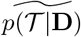—is shown in the upper right corner. We generated the plots using matplotlib Version 2.0.0 (Hunter, 2007).

To assess the effect of varying the number of demographic comparisons we repeated the simulations and analyses under Condition ***V***1, but with six demographic comparisons rather than three. Likewise, to assess the effect of varying the number of individuals sampled from each population, we repeated the simulations and analyses under Condition ***V***1, but with 20 individuals sampled per population (40 sampled genomes) rather than 10 (20 genomes).

#### 3.1.2 Simulations to assess sensitivity to prior assumptions

In the validation analyses above, the prior distributions used in analyses matched the true underlying distributions under which the data were generated. While this is an important first step when validating a Bayesian method and exploring its behavior under ideal conditions, this is unrealistic for real-world applications where our priors are always wrong and usually much more diffuse to express ignorance about the timing of past evolutionary events and historical effective population sizes. Also, having the priors match the true distributions effectively limits how extreme the simulating distributions can be. For example, the simulation condition ***V***1 above, where the distribution on the effective size of the ancestral population is sharply peaked at 0.25 (i.e., a four-fold population expansion), becomes a very informative prior distribution when analyzing the simulated data; more informative than is practical for most empirical applications of the method. Accordingly, we also analyzed data under conditions where the prior distributions are more diffuse than those under which the data were simulated. This allows us to (1) see how sensitive the method is to prior misspecification, (2) determine to what degree the results under conditions like ***V***1 are influenced by the sharply informative prior on the ancestral population size, and (3) explore more extreme simulation conditions of population expansions and contractions (conditions that would be unrealistic for a prior distribution).

We used the same distribution on event times and descendant effective population sizes as for Conditions ***V***1– ***V***5 above. For the relative effective size of the ancestral population 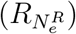, we chose four distributions under which to simulate data (***S***1–***S***4, Table 2) that are sharply peaked on four-fold and ten-fold population expansions and contractions. We simulated 500 data sets under each of these four conditions and then analyzed them under diffuse prior distributions. We chose the prior distributions to reflect realistic amounts of prior uncertainty about the timing of demographic changes and past and present effective population sizes when analyzing empirical data. Note, Conditions ***V***1 and ***S***1 share the same simulating distributions, which allows us to compare results to determine how much the strongly informative prior on the ancestral population size affected inference.

For comparison, we also repeated simulations and analyses under Conditions ***V***1 and ***S***1, except with three *divergence* comparisons. For these divergence comparisons, we simulated 10 sampled genomes per population to match the same total number of samples per comparison (20) as the demographic simulations.

#### 3.1.3 Simulating a mix of divergence and demographic comparisons

To explore how well our method can infer a mix of shared demographic changes and divergence times, we simulated 500 data sets comprised of 6 comparisons: 3 demographic comparisons and 3 divergence comparisons. To ensure the same amount of data across comparisons, we simulated 20 sampled genomes (10 diploid individuals) from each comparison (i.e., 10 genomes from each population of each divergence comparison). We used the same simulation conditions described above for ***V***2, and specified these same distributions as priors when analyzing all of the simulated data sets.

#### 3.1.4 Simulating linked sites

Our model assumes each character is effectively unlinked. To assess the effect of violating this assumption, we simulated data sets comprising 5000 100-base-pair loci (500,000 total characters). All 100 characters from each locus evolved along the same gene tree that is independent (conditional on the population history) from all other loci. The distributions on parameters were the same as the conditions described for ***V***1 above. These same distributions were used as priors when analyzing the simulated data sets.

### 3.2 Empirical application to stickleback data

#### 3.2.1 Assembly of loci

We assembled the publicly available RADseq data collected by Hohenlohe et al. (2010) from five populations of threespine sticklebacks (*Gasterosteus aculeatus*) from south-central Alaska. After downloading the reads mapped to the stickleback genome by Hohenlohe et al. (2010) from Dryad (doi:10.5061/dryad.b6vh6), we assembled reference guided alignments of loci in Stacks v1.48 Catchen et al. (2013) with a minimum read depth of 3 identical reads per locus within each individual and the bounded single-nucleotide polymorphism (SNP) model with error bounds between 0.001 and 0.01. To maximize the number of loci and minimize paralogy, we assembled each population separately; because `ecoevolity` models each population separately (Figure 1), the characters do not need to be orthologous across populations, only within them.

#### 3.2.2 Inferring shared demographic changes with ecoevolity

When analyzing the stickleback data with `ecoevolity`, we used a value for the concentration parameter of the Dirichlet process that corresponds to a mean number of three events (*α* = 2.22543). We used the following prior distributions on the timing of events and effective sizes of populations: *τ* ∼ Exponential(mean = 0.001), 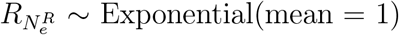, and 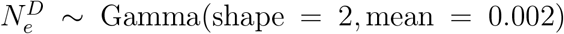. To assess the sensitivity of the results to these prior assumptions, we also analyzed the data under two additional priors on the concentration parameter, event times, and relative effective population size of the ancestral population:

- *α* = 13 (half of prior probability on 5 events)
- *α* = 0.3725 (half of prior probability on 1 event)
- *τ* ∼ Exponential(mean = 0.0005)
- *τ* ∼ Exponential(mean = 0.01)
- 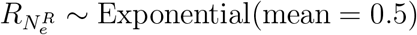
- 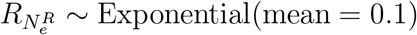

For each prior setting, we ran 10 MCMC chains for 150,000 generations, sampling every 100 generations; we did this using all the sites in the assembled stickleback loci and only variable sites (i.e., SNPs). To assess convergence and mixing of the chains, we calculated the PSRF (Brooks and Gelman, 1998) and ESS (Gong and Flegal, 2016) of all continuous parameters and the log likelihood using the `pyco-sumchains` tool of `pycoevolity` (Version 0.1.2 Commit 89d90a1). We also visually inspected the sampled log likelihood and parameter values over generations with the program Tracer (Version 1.6; Rambaut et al., 2014). The MCMC chains for all analyses converged almost immediately; we conservatively removed the first 101 samples from each chain, resulting in 14,000 samples from the posterior (1400 samples from 10 chains) for each analysis.

## 4 Results & Discussion

### 4.1 Analyses of simulated data

Despite our attempt to capture a mix of favorable and challenging parameter values in our initial simulation conditions (Table S1), estimates of the timing and sharing of demographic events were quite poor across all the simulation conditions we initially explored (see Supporting Information; Figures S1–S4). Even after we tried selecting simulation conditions that are more favorable for identifying the event times, estimates of the timing and sharing of demographic events remain quite poor (Figures 2 and 3). Under the recent (but not too recent) 4-fold population-size increase (on average) scenario, we do see better estimates of the times of demographic change (***V***1; top row of Figure 2), but the ability to identify the correct number of events and the assignment of the populations to those events remains quite poor; the correct model is preferred only 57% of the time, and the median posterior probability of the correct model is only 0.42 (top row of Figure 3). Under the most extreme population retraction scenario (***V***3; 4-fold, on average), the correct model is preferred only 40% of the time, and the median posterior probability of the correct model is only 0.26 (middle row of Figure 3). Estimates are especially poor when using only variable characters (second versus third column of Figures 2 and 3), so we focus on the results using all characters. We also see worse estimates of population sizes when excluding invariant characters (Figures S5 and S6).

Under the “worst-case” scenario of little population-size change (***V***5; bottom row of Figures 2 and 3), our method is unable to identify the timing or model of demographic change. As expected, under these conditions our method returns the prior on the timing of events (bottom row of Figure 2) and always prefers either a model with a single, shared demographic event (model “000”) or independent demographic changes (model “012”; bottom row of Figure 3). This is expected behavior, because there is essentially no information in the data about the timing of demographic changes, and a Dirichlet process with a mean of two demographic events, puts approximately 0.24 of the prior probability on the models with one and three events, and 0.515 prior probability on the three models with two events (approximately 0.17 each). Thus, with little information, the method samples from the prior distribution on the timing of events, and randomly prefers one of the two models with larger (and equal) prior probability.

Doubling the number of individuals sampled per population to 20 had very little effect on the results (Figure S7). Likewise, doubling the number of demographic comparisons to six had no effect on the accuracy or precision of estimating the timing of demographic changes or effective population sizes (Rows 1, 3, and 4 of Figure S8 and Figure S9). The ability to infer the correct number of demographic events, and assignment of populations to the events 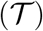, is much worse when there are six comparisons (Row 2 of Figure S8), which is not surprising given that the number of possible assignments of populations to events is 203 for six comparisons, compared to only five for three comparisons (Bell, 1934). We also see that the accuracy and precision of estimates of the timing of a demographic change event do not increase with the number of populations that share the event (Figure S9). This makes sense for two reasons: (1) it is difficult to correctly identify the sharing of demographic events among populations (Row 2 of Figure S8), and (2) Oaks (2019) and Oaks et al. (2019b) showed that the amount of information about the timing of events plateaus quickly as the number of characters increases. Thus, given 500,000 characters from each population, little information is to be gained about the timing of the demographic change, even if the method can correctly identify that several populations shared the same event.

The 95% credible intervals of all the parameters cover the true value approximately 95% of the time (Figures 2, S5, and S6). Given that our priors match the underlying distributions that generated the data, this coverage behavior is expected, and is an important validation of our implementation of the model and corresponding MCMC algorithms. The average run time of `ecoevolity` was approximately 21 and 42 minutes when analyzing three and six demographic comparisons, respectively. Analyses were run on a variety of hardware configurations, but most were run on 2.3GHz Intel Xeon CPU processors (E5-2650 v3).

#### 4.1.1 Sensitivity to prior assumptions

Above, we observe the best estimates of the timing and sharing of demographic events under the narrowest distribution on the relative effective size of the ancestral population (***V***1; top row of Figures 2 and 3), which was used to both simulate the data and as the prior when those data were analyzed. Thus, the improved behavior could be due to this narrow prior distribution that is unrealistically informative for most empirical studies, for which there is usually little *a priori* information about past population sizes. When we analyze data under more realistic, diffuse priors, estimates of the timing and sharing of *demographic* events deteriorate, whereas estimates of the timing and sharing of *divergence* events remain robust (Figures 4 and 5). Specifically, the precision of time estimates of demographic changes decreases substantially under the diffuse priors (top two rows of Figure 4), whereas the precision of the divergence-time estimates is high and largely unchanged under the diffuse priors (bottom two rows of Figure 4). We see the same patterns in the estimates of population sizes (Figures S10 and S11).

**Figure 4.**
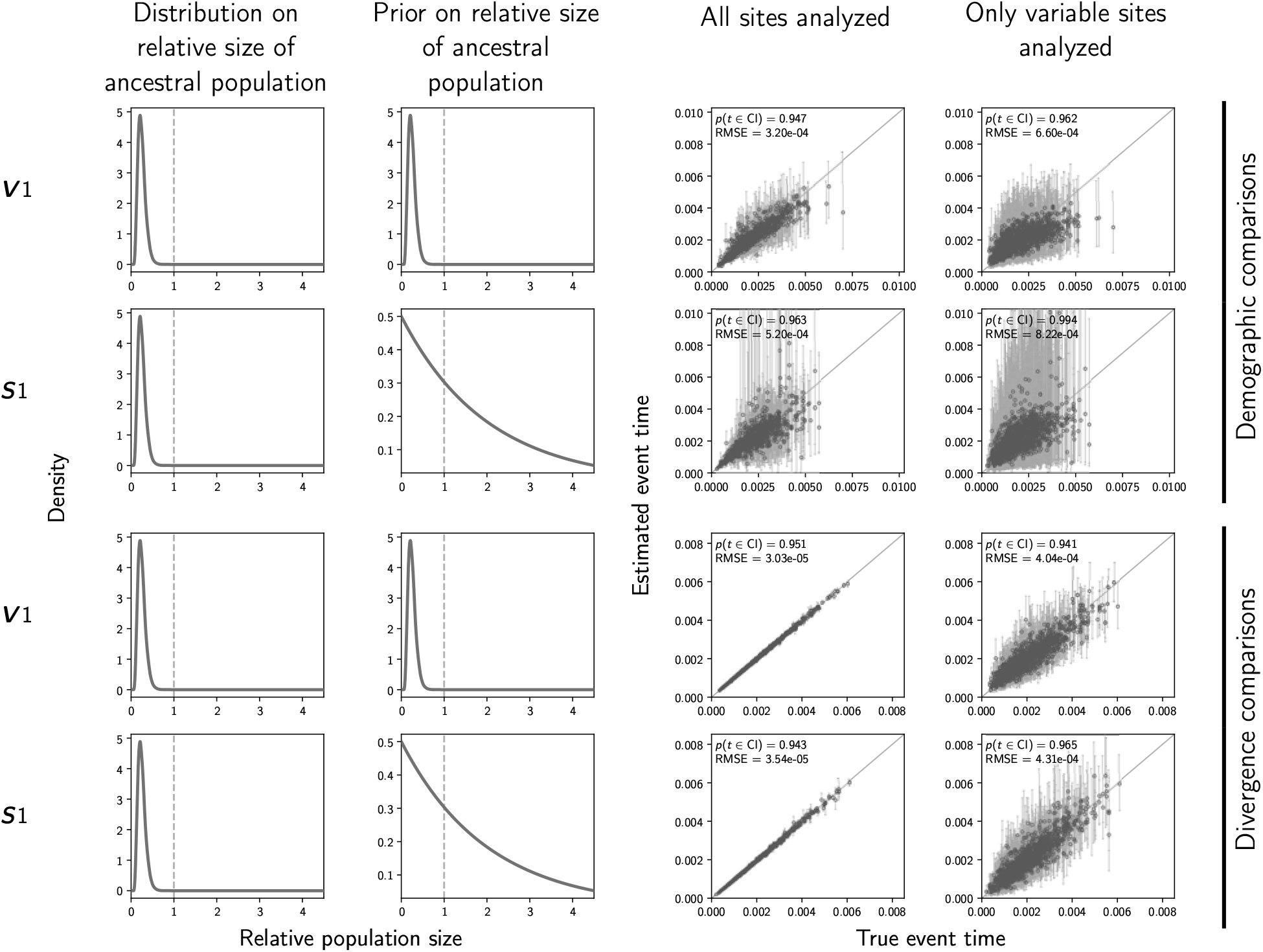
The accuracy and precision of time estimates of demographic changes (top two rows) versus divergences (bottom two rows) when the priors are correct (first and third rows) versus when the priors are diffuse (second and fourth rows). Time is measured in units of expected subsitutions per site. The first and second columns of plots show the distribution on the relative effective size of the ancestral population for simulating the data (Column 1) and for the prior when analyzing the simulated data (Column 2). The third and fourth columns of plots show true versus estimated values when using all characters (Column 3) or only variable characters (Column 4). Each plotted circle and associated error bars represent the posterior mean and 95% credible interval. Estimates for which the potential-scale reduction factor was greater than 1.2 (Brooks and Gelman, 1998) are highlighted in orange. Each plot consists of 1500 estimates—500 simulated data sets, each with three demographic comparisons (Rows 1–2) or divergence comparisons (Rows 3–4). For each plot, the root-mean-square error (RMSE) and the proportion of estimates for which the 95% credible interval contained the true value—*p*(*t* ∈ CI)—is given. The first row of plots are repeated from Figure 2 for comparison. We generated the plots using matplotlib Version 2.0.0 (Hunter, 2007).

**Figure 5.**
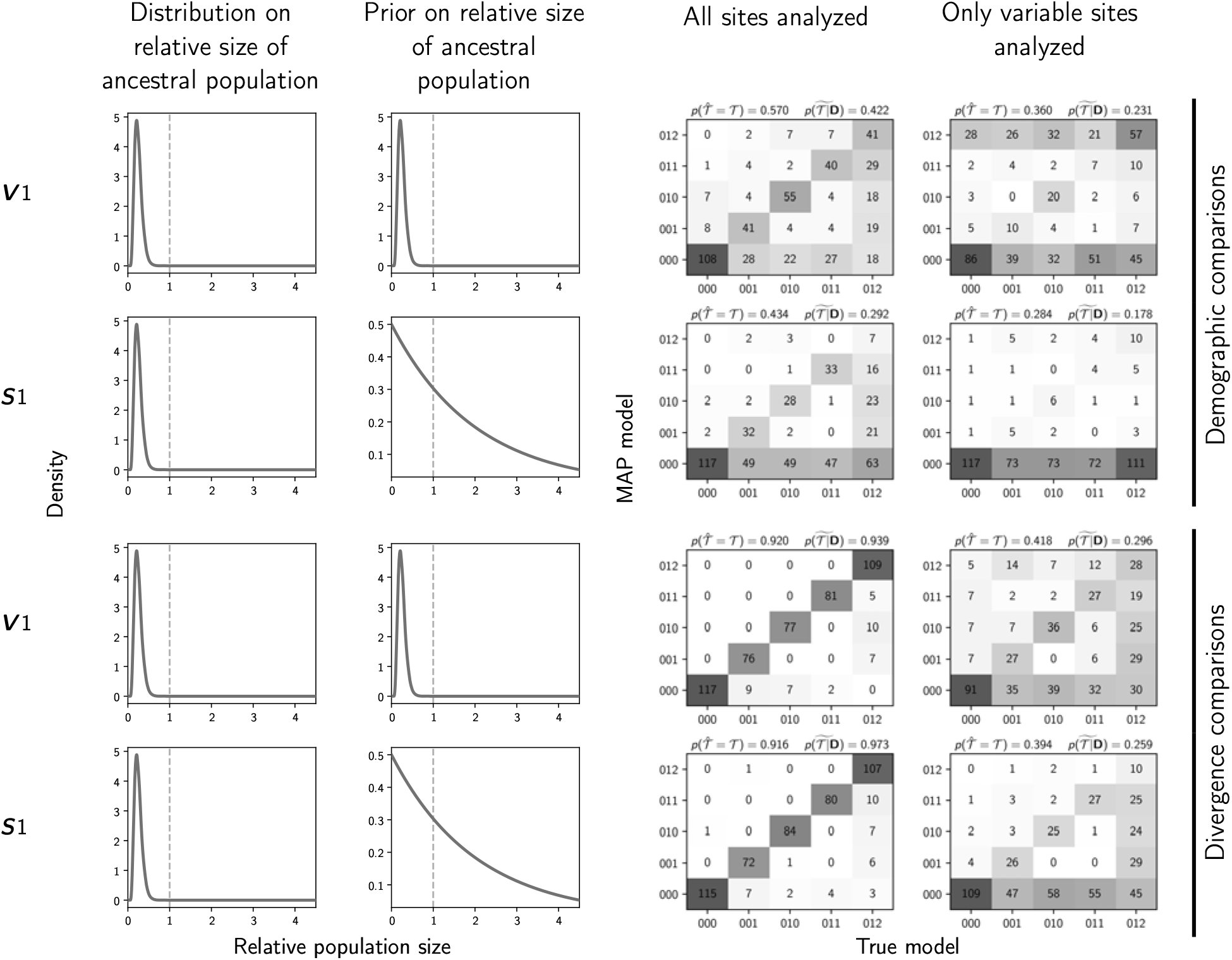
The performance of estimating the model of demographic changes (top two rows) versus model of divergences (bottom two rows) when the priors are correct (first and third rows) versus when the priors are diffuse (second and fourth rows). The first and second columns of plots show the distribution on the relative effective size of the ancestral population for simulating the data (Column 1) and for the prior when analyzing the simulated data (Column 2). The third and fourth columns of plots show true versus estimated models when using all characters (Column 3) or only variable characters (Column 4). Each plot shows the results of the analyses of 500 simulated data sets, each with three demographic comparisons (Rows 1–2) or divergence comparisons (Rows 3–4); the number of data sets that fall within each possible cell of true versus estimated model is shown, and cells with more data sets are shaded darker. Each model is represented along the plot axes by three integers that indicate the event category of each comparison (e.g., 011 represents the model in which the second and third comparison share the same event time that is distinct from the first). The estimates are based on the model with the maximum *a posteriori* probability (MAP). For each plot, the proportion of data sets for which the MAP model matched the true model—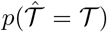—is shown in the upper left corner, and the median posterior probability of the correct model across all data sets—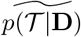—is shown in the upper right corner. The first row of plots are repeated from Figure 3 for comparison. We generated the plots using matplotlib Version 2.0.0 (Hunter, 2007).

Furthermore, under the diffuse priors, the probability of inferring the correct model of demographic events decreases from 0.57 to 0.434 when all characters are used, and from 0.36 to 0.284 when only variable characters are used (top two rows of Figure 5). The median posterior probability of the correct model also decreases from 0.422 to 0.292 when all characters are used, and from 0.231 to 0.178 when only variable characters are used (top two rows of Figure 5). Most importantly, we see a strong bias toward underestimating the number of events under the more realistic diffuse priors (top two rows of Figure 5). In comparison, the inference of shared divergence times is much more accurate, precise, and robust to the diffuse priors (bottom two rows of Figure 5). When all characters are used, under both the correct and diffuse priors, the correct divergence model is preferred over 91% of the time, and the median posterior probability of the correct model is over 0.93.

Results are very similar whether the distribution on the ancestral population size is peaked around a four-fold population expansion or contraction (Conditions ***S***1 and ***S***2; top two rows of Figures 6, 7, S12, and S13). Likewise, even when population expansions and contractions are 10-fold, the ability to infer the timing and sharing of these events remains poor (Conditions ***S***3 and ***S***4; bottom two rows of Figures 6 and 7). This is not surprising when reflecting on the first principles of this inference problem. While it may seem intuitive that more dramatic changes in the rate of coalescence should be easier to detect, such large changes will cause fewer lineages to coalesce after (in the case of a dramatic population expansion) or before (in the case of a dramatic population contraction) the change in population size. This reduces the information about the rate of coalescence on one side of the demographic change and thus the magnitude and timing of the change in effective population size. Thus, the gain in information in the data is expected to plateau (and even decrease, as we see under the most severe bottleneck Condition ***S***4 in Figure 7) as the magnitude of the change in effective population size increases.

**Figure 6.**
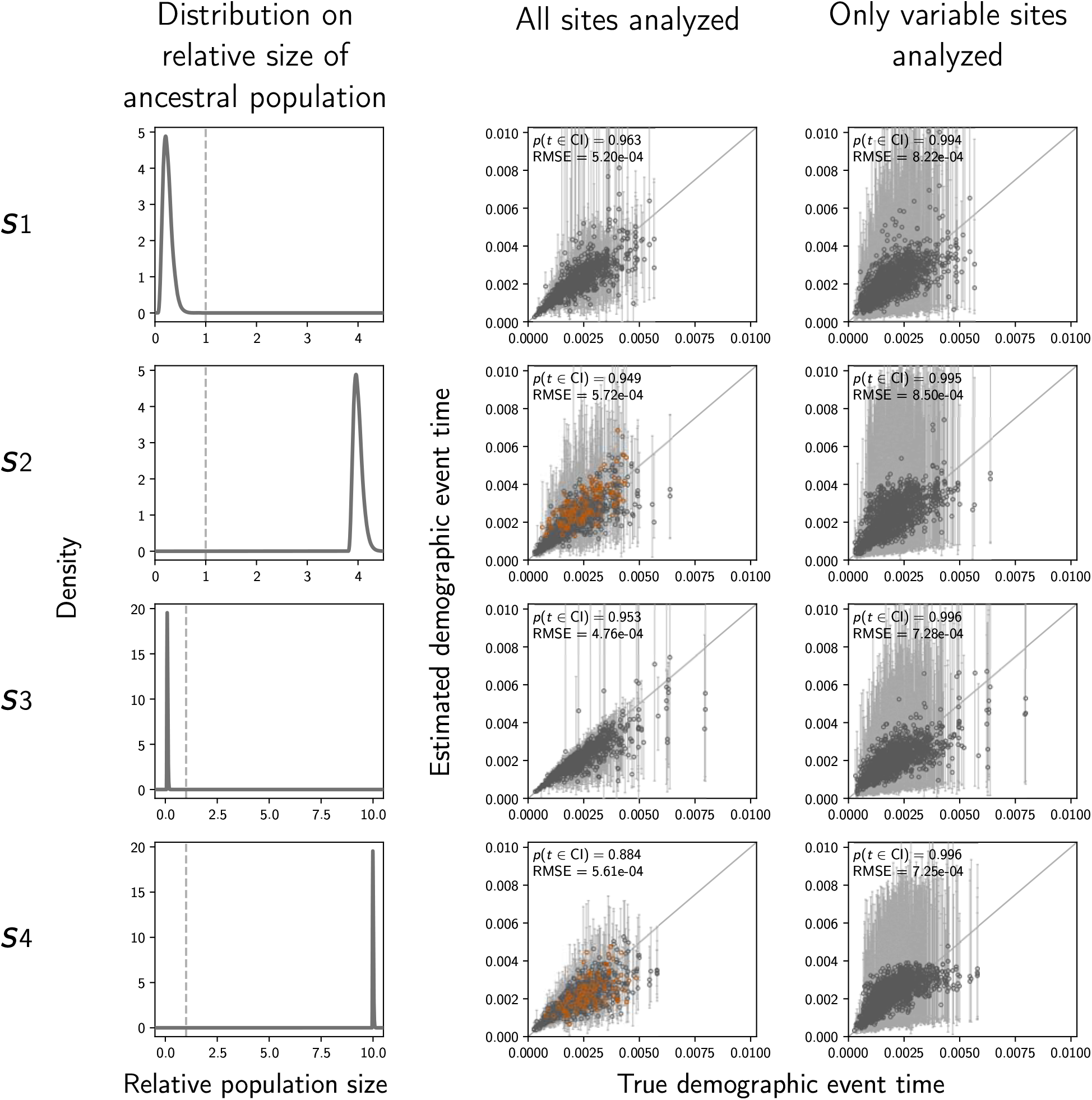
The accuracy and precision of time estimates of demographic changes when the prior distributions are diffuse (Conditions ***S***1–***S***4; Table 2). Time is measured in units of expected subsitutions per site. The first column of plots shows the distribution on the relative effective size of the ancestral population under which the data were simulated, and the second and third columns of plots show true versus estimated values when using all characters (Column 2) or only variable characters (Column 3). Each plotted circle and associated error bars represent the posterior mean and 95% credible interval. Estimates for which the potential-scale reduction factor was greater than 1.2 (Brooks and Gelman, 1998) are highlighted in orange. Each plot consists of 1500 estimates—500 simulated data sets, each with three demographic comparisons. For each plot, the root-mean-square error (RMSE) and the proportion of estimates for which the 95% credible interval contained the true value—*p*(*t* ∈ CI)—is given. The first row of plots are repeated from Figure 4 for comparison. We generated the plots using matplotlib Version 2.0.0 (Hunter, 2007).

**Figure 7.**
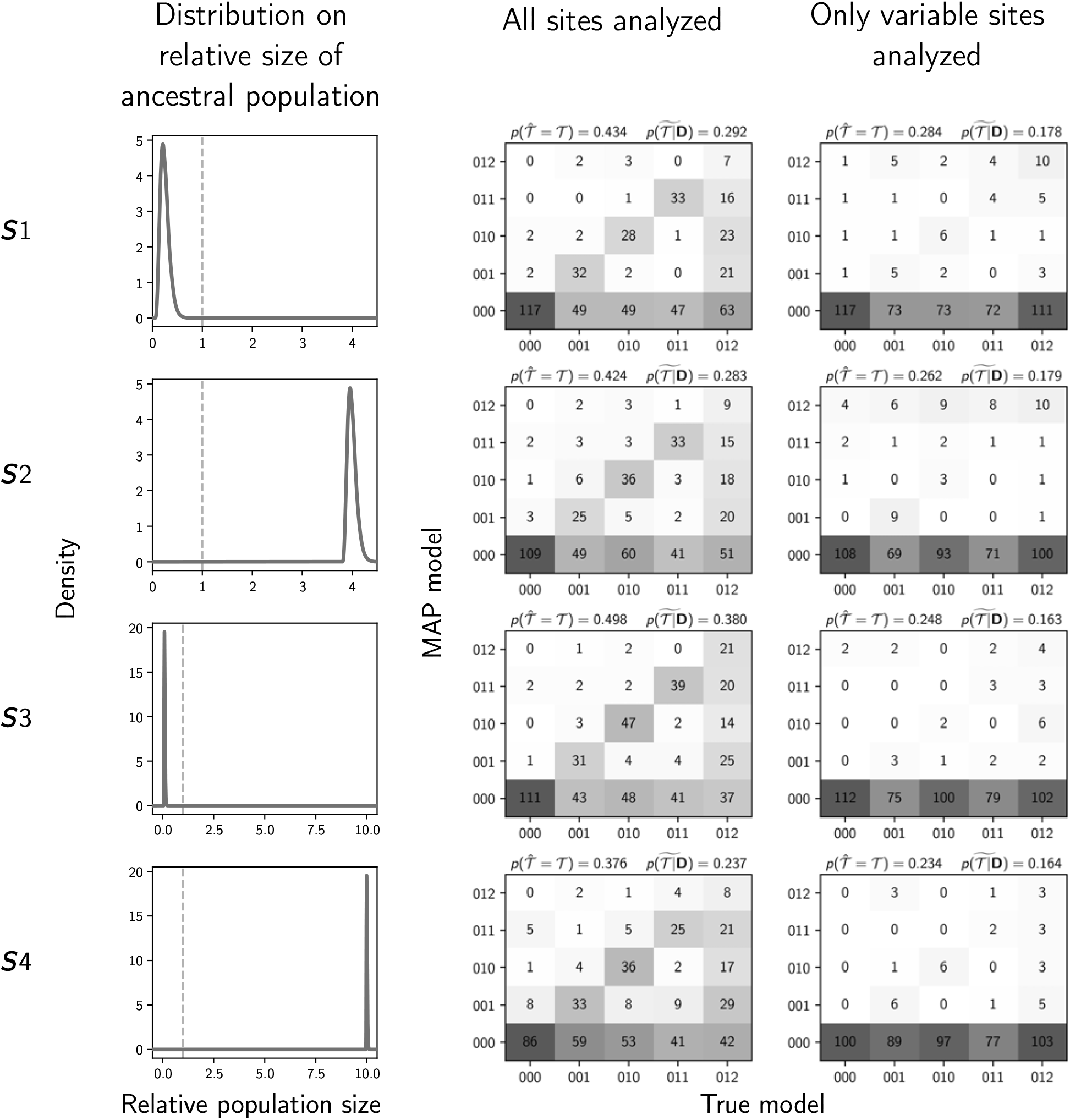
The performance of estimating the model of demographic changes when the prior distributions are diffuse (Conditions ***S***1–***S***4; Table 2). The first column of plots shows the distribution on the relative effective size of the ancestral population under which the data were simulated, and the second and third columns of plots show true versus estimated models when using all characters (Column 2) or only variable characters (Column 3). Each plot consists of 1500 estimates—500 simulated data sets, each with three demographic comparisons; the number of data sets that fall within each possible cell of true versus estimated model is shown, and cells with more data sets are shaded darker. Each model is represented along the plot axes by three integers that indicate the event category of each comparison (e.g., 011 represents the model in which the second and third comparison share the same event time that is distinct from the first). The estimates are based on the model with the maximum *a posteriori* probability (MAP). For each plot, the proportion of data sets for which the MAP model matched the true model—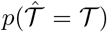—is shown in the upper left corner, and the median posterior probability of the correct model across all data sets—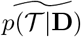—is shown in the upper right corner. The first row of plots are repeated from Figure 4 for comparison. We generated the plots using matplotlib Version 2.0.0 (Hunter, 2007).

#### 4.1.2 Inferring a mix of shared divergences and demographic changes

When demographic and divergence comparisons are analyzed separately, the performance of estimating the timing and sharing of demographic changes and divergences is dramatically different, with the latter being much more accurate and precise than the former (e.g., see Figures 4 and 5). One might hope that if we analyze a mix of demographic and divergence comparisons, the informativeness of the divergence times can help “anchor” and improve the estimates of shared demographic changes. However, our results from simulating data sets comprising a mix of three demographic and three divergence comparisons rule out this possibility. When analyzing a mix of demographic and divergence comparisons, the ability to infer the timing and sharing of demographic changes remains poor, whereas estimates of shared divergences remain accurate and precise (Figure 8). The estimates of the timing and sharing of demographic events are nearly identical to when we simulated and analyzed only three demographic comparisons under the same distributions on event times and population sizes (Condition ***V***2; compare left column of Figure 8 to the second row of Figures 2 and 3). The same is true for the estimates of population sizes (Figure S14). Thus, there does not appear to be any mechanism by which the more informative divergence-time estimates “rescue” the estimates of the timing and sharing of the demographic changes.

**Figure 8.**
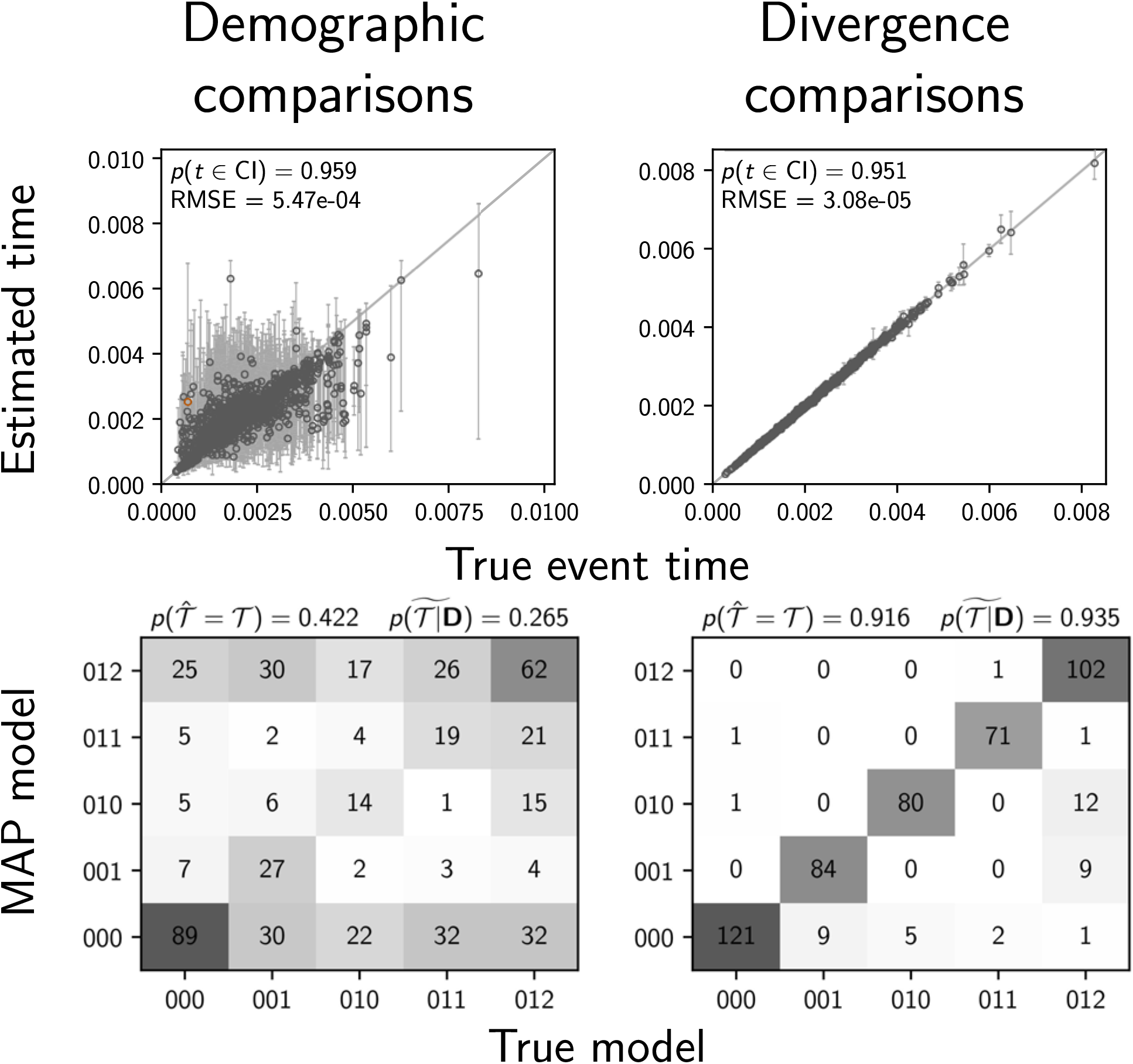
Results of analyses of 500 data sets simulated with six comparisons comprising a mix of three populations that experienced a demographic change and three pairs of populations that diverged. The performance of estimating the timing (top row) and sharing (bottom row) of events are shown separately for the three populations that experienced a demographic change (left column) and the three pairs of populations that diverged (right column). The plots of the demographic comparisons (left column) are comparable to the second column of Figures 2 and 3; the same priors on event times and ancestral population size were used. Time estimates for which the potential-scale reduction factor was greater than 1.2 (Brooks and Gelman, 1998) are highlighted in orange. We generated the plots using matplotlib Version 2.0.0 (Hunter, 2007).

#### 4.1.3 The effect of linked sites

Most reduced-representation genomic datasets are comprised of loci of contiguous, linked nucleotides. Thus, when using the method presented here that assumes each character is effectively unlinked, one either has to violate this assumption, or discard all but (at most) one site per locus. Given that all the results above indicate better estimates when all characters are used (compared to using only variable characters), we simulated linked sites to determine which strategy is better: analyzing all linked sites and violating the assumption of unlinked characters, or discarding all but (at most) one variable character per locus.

The results are almost identical to when all the sites were unlinked (compare first row of Figures 2 and 3 to the top two rows of Figure S15, and the first row of Figures S5 and S6 to the bottom two rows of Figure S15). Thus, violating the assumption of unlinked sites has little effect on the estimation of the timing and sharing of demographic changes or the effective population sizes. This is consistent with the findings of Oaks (2019) and Oaks et al. (2019b) that linked sites had little impact on the estimation of shared divergence times. These results suggest that analyzing all of the sites in loci assembled from reduced-representation genomic libraries (e.g., sequence-capture or RADseq loci) is a better strategy than excluding sites to avoid violating the assumption of unlinked characters.

### 4.2 Reassessing the co-expansion of stickleback populations

Using an ABC analog to the model of shared demographic changes developed here, Xue and Hickerson (2015) found very strong support (0.99 posterior probability) that five populations of threespine sticklebacks (*Gasterosteus aculeatus*) from south-central Alaska recently co-expanded. This inference was based on the publicly available RADseq data collected by Hohenlohe et al. (2010). We re-assembled and analyzed these data under our full-likelihood Bayesian framework, both using all sites from assembled loci and only variable sites (i.e., SNPs).

Stacks produced a concatenated alignment with 2,115,588, 2,166,215, 2,081,863, 2,059,650, and 2,237,438 total sites, of which 118,462, 89,968, 97,557, 139,058, and 103,271 were variable for the Bear Paw Lake, Boot Lake, Mud Lake, Rabbit Slough, and Resurrection Bay stickleback populations respectively. When analyzing all sites from the assembled stickleback RADseq data, we find strong support for five independent population expansions (no shared demographic events; Figure 9). In sharp contrast, when analyzing only SNPs, we find support for a single, shared, extremely recent population expansion (Figure 9). These results are relatively robust to a broad range of prior assumptions (Figures S16–24). The support for a single, shared event is consistent with the results from our simulations using diffuse priors and only including SNPs, which showed consistent, spurious support for a single event (Row 2 of Figure 5 and Figure 7).

**Figure 9.**
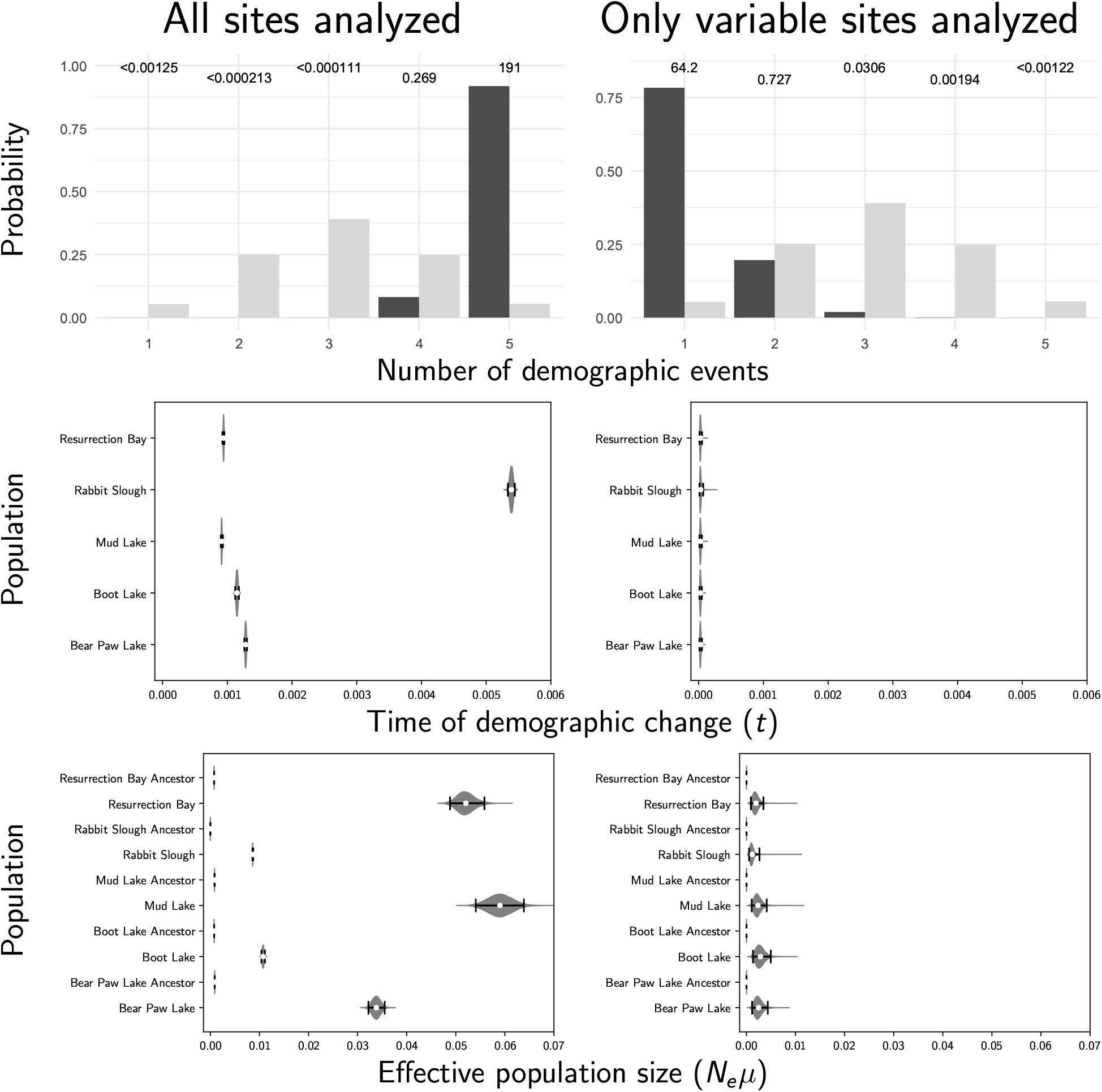
Estimates of the number (Row 1), timing (Row 2), and magnitude (Row 3) of demographic events across five stickleback populations, when using all sites (left column) or only variable sites (right column). We used an exponentially distributed prior with a mean of 0.001 on event times, an exponentially distributed prior with a mean of 1 on the relative ancestral effective population size, and a gamma-distributed prior (shape = 2, mean = 0.002) on the descendant population sizes. For the number of events (Row 1), the light and dark bars represent the prior and posterior probabilities, respectively. The numbers above the bars are Bayes factors (the ratio of posterior-to-prior odds) comparing each number of events to all other numbers of events. Time (Row 2) is in units of expected subsitutions per site. For the violin plots, each plotted circle and associated error bars represent the posterior mean and 95% credible interval. Bar graphs were generated with ggplot2 Version 2.2.1 (Wickham, 2009); violin plots were generated with matplotlib Version 2.0.0 (Hunter, 2007).

When using only SNPs, estimates of the timing of the single, shared demographic event from the stickleback data are essentially at the minimum of zero (Figure 9), suggesting that there is little information about the timing of any demographic changes in the SNP data alone. This is consistent with results of Xue and Hickerson (2015) where the single, shared event was also estimated to have occurred at the minimum (1000 generations) of their uniform prior on the timing of demographic changes. In light of our simulation results, the support for a single event based solely on SNPs, seen here and in Xue and Hickerson (2015), is likely caused by a combination of (1) misspecified priors, and (2) the lack of information about demographic history when invariant characters are discarded. By saying the priors were misspecified, we mean that the prior distributions do not match the true distributions underlying the generation of the data, not that the priors were poorly chosen. Our estimates using all of the sites in the stickleback RADseq loci should be the most accurate, according to our results from simulated data. However, the unifying theme from our simulations is that all estimates of shared demographic events tend to be poor and should be treated with a lot of skepticism.

### 4.3 Biological realism of our model of shared demographic changes

The model of shared population-size changes we present above, and used in previous research (Chan et al., 2014; Xue and Hickerson, 2015; Gehara et al., 2017; Xue and Hickerson, 2015), is quite unrealistic in number of ways. Modeling the demographic history of a population with a single, instantaneous change in population size does not reflect the continuous and complex demographic changes most populations of organisms experience through time. However, this simple model is correct in our simulated data, and yet our method struggles to accurately infer the timing and sharing of these single, dramatic, instantaneous changes in effective population size. Incorporating more demographic realism into the model will introduce more variation and thus make the inference problem even more difficult. Thus, until inference of shared events under overly simplistic demographic models can be improved, it does not seem wise to introduce more complexity.

Also, we expect most processes that cause shared divergences and/or demographic changes across species will affect multiple species with some amount of temporal variation. Thus, our model of simultaneous evolutionary events that affect multiple species at the same instant is not biologically plausible. If this lack of realism is problematic, it should cause the method to overestimate the number of events by misidentifying the temporal variation among species affected by the same process as being the result of multiple events. However, what we see here (e.g., Figure 7) and what has been shown previously (Oaks et al., 2013, 2014; Oaks, 2014, 2019; Oaks et al., 2019b) is the opposite; even when we model shared events as simultaneous, methods tend to underestimate (and almost never overestimate) the number of events. We do see overestimates when there is little information in the data and the posterior largely reflects the prior (e.g., bottom two rows of Figure 3). However, this is only true when the prior distributions match the true underlying distributions that generated the data, and these overestimates would be easy to identify in practice by testing for prior sensitivity and noticing that the posterior probabilities of event models are similar to the prior probabilities (i.e., small Bayes factors). Furthermore, Oaks et al. (2019b) showed that even with millions of bases of genomic data from pairs of gecko populations, `ecoevolity` was only able to detect differences in divergence times between comparisons greater than several thousand years. Thus, it seems unlikely that over-estimating the number of events among taxa (i.e., estimating temporal independence of comparisons that shared the same historical process) is a real problem for these types of inferences.

Previous researchers (Overcast et al., 2017; Gehara et al., 2017; Xue and Hickerson, 2017) have attempted to introduce realism into these comparative models by allowing temporal variation among species affected by the same event, by assuming that processes of diversification and demographic change are temporally overdispersed. However, allowing temporal variation within events will only increase the tendency of these methods to underestimate the number of events (i.e., the within-event temporal variation makes it “easier” to assign comparisons to the same event). More fundamentally, it seems odd to assume *a priori* that processes that cause shared evolutionary responses would be somehow conveniently staggered over evolutionary timescales (overdispersed); this seems like something we would want to estimate from the data.

### 4.4 Comparison to previous models of shared demographic changes

Our method is the first that we know of that is generalized to infer an arbitrary mix of shared times of divergence and changes in population size. However, if we focus only on changes in population size, the models underlying the ABC methods of Chan et al. (2014), Xue and Hickerson (2015), and Gehara et al. (2017) share many similarities with the model we introduced above. These models, like ours, allow the effective population sizes before and after the time of the demographic change to vary (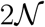 free parameters), however, they assume all populations experienced an expansion. The models of Chan et al. (2014), Xue and Hickerson (2015), and Gehara et al. (2017) also assume there was at most one shared demographic event; each comparison can either be assigned to this event or have an independent time of demographic change. Xue and Hickerson (2017) relaxed these constraints by allowing population contractions and expansions and allowing any number of demographic events and assignments of populations to those events, like we do here. All previous approaches, like ours, model variation in gene trees using the coalescent. They also assume an infinite-sites model of character evolution along gene trees, whereas our approach uses a finite-sites model. Gehara et al. (2017) and Xue and Hickerson (2017) allow the investigator to assume that the processes that cause demographic changes are temporally overdispersed (i.e., separated in time by “buffers”). We do not explore this temporal staggering of events here because this is a pattern we would like to infer from data rather than impose *a priori*. Furthermore, creating temporal “buffers” around events will exacerbate the tendency to over-cluster comparisons (i.e., underestimate the number of events).

The biggest difference between previous approaches and ours is how the data are used. Chan et al. (2014) and Gehara et al. (2017) reduce aligned sequences into a set of population genetic summary statistics. Xue and Hickerson (2015) and Xue and Hickerson (2017) reduce SNPs into a site-frequency spectrum (SFS) that is aggregated across the populations being compared. Both of these approaches to summarizing the data result in information about the model being lost (i.e., the summary statistics used for inference are insufficient). By using the mathematical work of Bryant et al. (2012), our method is able to calculate the likelihood of the population histories of the comparisons directly from the counts of character patterns from each population, while integrating over all possible gene trees under a coalescent model and all possible mutational histories along those gene trees under a finite-sites model of character evolution. Not only does this allow our approach to leverage all of the information in the data, but it does so efficiently; when analyzing four divergence comparisons, Oaks (2019) found this approach requires approximately 9340 times less computing time than using ABC. Also, calculating the likelihood of the model from each character pattern naturally accommodates missing data (Oaks, 2019). In contrast, there is no straightforward way of accounting for missing data when summarizing genetic data into SFS and other population genetic summary statistics (Hahn, 2018).

## 5 Conclusions

There is a narrow temporal window within which we can reasonably estimate the time of a demographic change. The width of this window is determined by how deep in the past the change occurred relative to the effective size of the population (i.e., in coalescent units). If too old or recent, there are too few coalescence events before or after the demographic change, respectively, to provide information about the effective size of the population. When we are careful to simulate data within this window, and the change in population size is large enough, we can estimate the time of the demographic changes reasonably well (e.g., see the top row of Figure 2). However, even under these favorable conditions, the ability to correctly infer the shared timing of demographic events among populations is quite limited (Figure 3). When only variable characters are analyzed (i.e., SNPs), estimates of the timing and sharing of demographic changes are consistently bad; we see this across all the conditions we simulated. Most alarmingly, when the priors are more diffuse than the distributions that generated the data, as will be true in most empirical applications, there is a strong bias toward estimating too few demographic events (i.e., over-clustering comparisons to demographic events; Row 2 of Figure 5), especially when only variable characters are analyzed. These results help explain the stark contrast we see in our results from the stickleback RADseq data when including versus excluding constant sites (Figure 9). These findings are in sharp contrast to estimating shared *divergence* times, which is much more accurate, precise, and robust to prior assumptions (Figures 4, 5 and 8; Oaks, 2019; Oaks et al., 2019b).

Given the poor estimates of co-demographic changes, even when all the information in the data are leveraged by a full-likelihood method, any inference of shared demographic changes should be treated with caution. However, there are potential ways that estimates of shared demographic events could be improved. For example, as discussed by Myers et al. (2008), modelling loci of contiguous, linked sites could help extract more information about past demographic changes. Longer loci can contain much more information about the lengths of branches in the gene tree, which are critically informative about the size of the population through time. This is evidenced by the extensive literature on powerful “skyline plot” and “phylodynamic” methods (Pybus et al., 2000; Strimmer and Pybus, 2001; Opgen-Rhein et al., 2005; Drummond et al., 2005; Heled and Drummond, 2008; Minin et al., 2008; Ho and Shapiro, 2011; Palacios and Minin, 2013, 2012; Stadler et al., 2013; Gill et al., 2013; Palacios et al., 2014; Lan et al., 2015; Karcher et al., 2016, 2017; Faulkner et al., 2018; Karcher et al., 2019). Obviously, the length of loci will be constrained by recombination. Nonetheless, with loci from across the genome, each with more information about the gene tree they evolved along (Speidel et al., 2019), perhaps more information can be captured about temporally clustered changes in the rate of coalescence across populations.

Another potential source of information could be captured by modelling recombination along large regions of chromosomes. By approximating the full coalescent process, many methods have been developed to model recombination in a computationally feasible manner (McVean and Cardin, 2005; Marjoram and Wall, 2006; Chen et al., 2009; Li and Durbin, 2011; Sheehan et al., 2013; Schiffels and Durbin, 2014; Rasmussen et al., 2014; Palacios et al., 2015). This could potentially leverage additional information from genomic data about the linkage patterns among sites along chromosomes.

The inference of shared evolutionary events could also stand to benefit from information about past environmental conditions, life history data about the taxa, and ecological data about how they interact. Modeling ecological aspects of the taxa and historical environmental conditions could provide important information about which comparisons are most likely to respond to environmental changes and when, and which taxa are likely to interact and influence each other’s demographic trajectories. While collecting these types of data and modelling these sorts of dynamics is challenging, approximate approaches can help to lead the way (He et al., 2013; Massatti and Knowles, 2016; Bemmels et al., 2016; Knowles and Massatti, 2017; Papadopoulou and Knowles, 2016). All of the these avenues are worth pursuing given the myriad historical processes that predict patterns of temporally clustered demographic changes across species.

## 6 Acknowledgments

We thank the members of the Phyletica Lab (the phyleticians) for helpful feedback on multiple drafts of this paper. This work was supported by funding provided to JRO from the National Science Foundation (NSF grant number DEB 1656004). NL was supported for a summer REU from NSF grant DEB 1656004 to JRO; NL also benefitted from an NSF REU award to Dr. Leslie Goertzen (grant number DBI 1560115). The computational work was made possible by the Auburn University (AU) Hopper Cluster supported by the AU Office of Information Technology and a grant of high-performance computing resources and technical support from the Alabama Supercomputer Authority. We thank Graham Stone, Michael Hickerson, and two anonymous reviewers for helpful feedback about this work. This paper is contribution number 899 of the Auburn University Museum of Natural History.

## 7 Data Accessibility

A detailed history of all aspects of this project was recorded in a version-controlled repository, which is publicly available at https://github.com/phyletica/ecoevolity-demog-experiments, and was archived on zenodo (https://doi.org/10.5281/zenodo.3319992; Oaks et al., 2019a).

## 8 Author Contributions

J.R.O. conceived the study. J.R.O. and N.L. designed and executed the simulation analyses. K.A.C. assembled the stickleback sequence data, and J.R.O. and K.A.C. analyzed those data. J.R.O. led the writing of the manuscript, with contributions from N.L. and K.A.C.

## Supporting Information

## 1 Methods

### 1.1 Initial simulation conditions

We initially simulated data under distributions we hoped comprised a mix of conditions that were favorable and challenging for estimating the timing and sharing of demographic changes. For these initial conditions, we simulated data sets with three populations that underwent a demographic change, under five different distributions on the relative effective size of the ancestral population (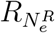; see Table S1 and left column of Figures S1 and S2), which ranged from having a mean 4-fold population-size increase (***PV***1) to a 2-fold decrease (***PV***3) and a “worst-case” scenario where there was essentially no population-size change in the history of the populations (***PV***5).

**Table S1.** Simulation and analysis conditions for preliminary validation analyses. The distributions from which parameter values were drawn for simulating data with simcoevolity are given for event times (*τ*), the relative effective size of the root (ancestral) population 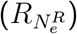, and the effective size of the descendant population 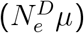, along with the prior distributions used for these parameters when the simulated data sets were analyzed with ecoevolity. When the latter is represented by a dash, this means the prior distribution matched the distribution under which the data were simulated. G(…) and E(…) represent gamma and exponential distributions, respectively, and the first number provided for the gamma distributions is the shape parameter.

| Label | Simulated distribution |  |  | Prior distribution |  |  |
| --- | --- | --- | --- | --- | --- | --- |
| | $\tau$ | $R_{N_e^R}$ | $N_e^D\mu$ | $\tau$ | $R_{N_e^R}$ | $N_e^D\mu$ |
| **Preliminary validation simulation conditions** |  |  |  |  |  |  |
| **PV1** | $E(\text{mean} = 0.01)$ | $G(10, \text{mean} = 0.25)$ | $G(5, \text{mean} = 0.002)$ | - | - | - |
| **PV2** | $E(\text{mean} = 0.01)$ | $G(10, \text{mean} = 0.5)$ | $G(5, \text{mean} = 0.002)$ | - | - | - |
| **PV3** | $E(\text{mean} = 0.01)$ | $G(10, \text{mean} = 2)$ | $G(5, \text{mean} = 0.002)$ | - | - | - |
| **PV4** | $E(\text{mean} = 0.01)$ | $G(10, \text{mean} = 1)$ | $G(5, \text{mean} = 0.002)$ | - | - | - |
| **PV5** | $E(\text{mean} = 0.01)$ | $G(100, \text{mean} = 1)$ | $G(5, \text{mean} = 0.002)$ | - | - | - |

For the mutation-scaled effective size of the descendant populations (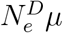; i.e., the population size after the demographic change), we used a gamma distribution with a shape of 5 and mean of 0.002 (Table S1). The timing of the demographic events was exponentially distributed with a mean of 0.01 substitutions per site. Taken together, the mean of the distribution on event times in units of 4*N_e_* generations is approximately 1.56. We chose this distribution in order to span times of demographic change from very recent (i.e., most gene lineages coalesce before the change) to old (i.e., most gene lineages coalesce after the change), which we assumed would include conditions under which the method performed both well and poorly. The assignment of the population-size change of the three simulated populations to 1, 2, or 3 demographic events was controlled by a Dirichlet process with a mean number of two events across the three populations. We generated 500 data sets under each of these five simulation conditions, all of which were analyzed using the same simulated distributions as priors.

## 2 Results

Despite our attempt to capture a mix of favorable and challenging parameter values, estimates of the timing (Figure S1) and sharing (Figure S2) of demographic events were quite poor across all the simulation conditions we initially explored. Under the “worst-case” scenario of very little population-size change (bottom row of Figures S1 and S2), our method is unable to identify the timing or model of demographic change. Under these conditions, our method returns the prior on the timing of events (bottom row of Figure S1) and almost always prefers either a model with a single, shared demographic event (model “000”) or independent demographic changes (model “012”; bottom row of Figure S2). This behavior is expected, because there is very little information in the data about the timing of demographic changes, and a Dirichlet process with a mean of 2.0 demographic events, puts approximately 0.24 of the prior probability on the models with one and three events, and 0.515 prior probability on the three models with two events (approximately 0.17 each). As a result, with little information, the method samples from the prior distribution on the timing of events, and prefers one of the two models with the largest (and equal) prior probability.

Under considerable changes in population size, the method only fared moderately better at estimating the timing of demographic events (top three rows of Figure S1). The ability to identify the model improved under these conditions, but the frequency at which the correct model was preferred only exceeded 50% for the large population expansions (top two rows of Figure S2). The median posterior support for the correct model was very small (less than 0.58) under all conditions. Under all simulation conditions, estimates of the timing and sharing of demographic events are better when using all characters, rather than only variable characters (second versus third column of Figures S1 and S2). Likewise, we see better estimates of effective population sizes when using the invariant characters (Figures S3 and S4).

We observed numerical problems when the time of the demographic change was either very recent or old relative to the effective size of the population following the change (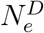; the descendant population). In such cases, either very few or almost all of the sampled gene copies coalesce after the demographic change, providing almost no information about the magnitude or timing of the population-size change. In these cases, the data are well-explained by a constant population size, which can be achieved by the model in three ways: (1) an expansion time of zero and an ancestral population size that matched the true population size, (2) an old expansion and a descendant population size that matched the true population size, or (3) an intermediate expansion time and both the ancestral and descendant sizes matched the true size. The true population size being matched in these modelling conditions is that of the descendant or ancestral population if the expansion was old or recent, respectively. This caused MCMC chains to converge to different regions of parameter space (highlighted in orange in Figure S1).

**Figure S1.**
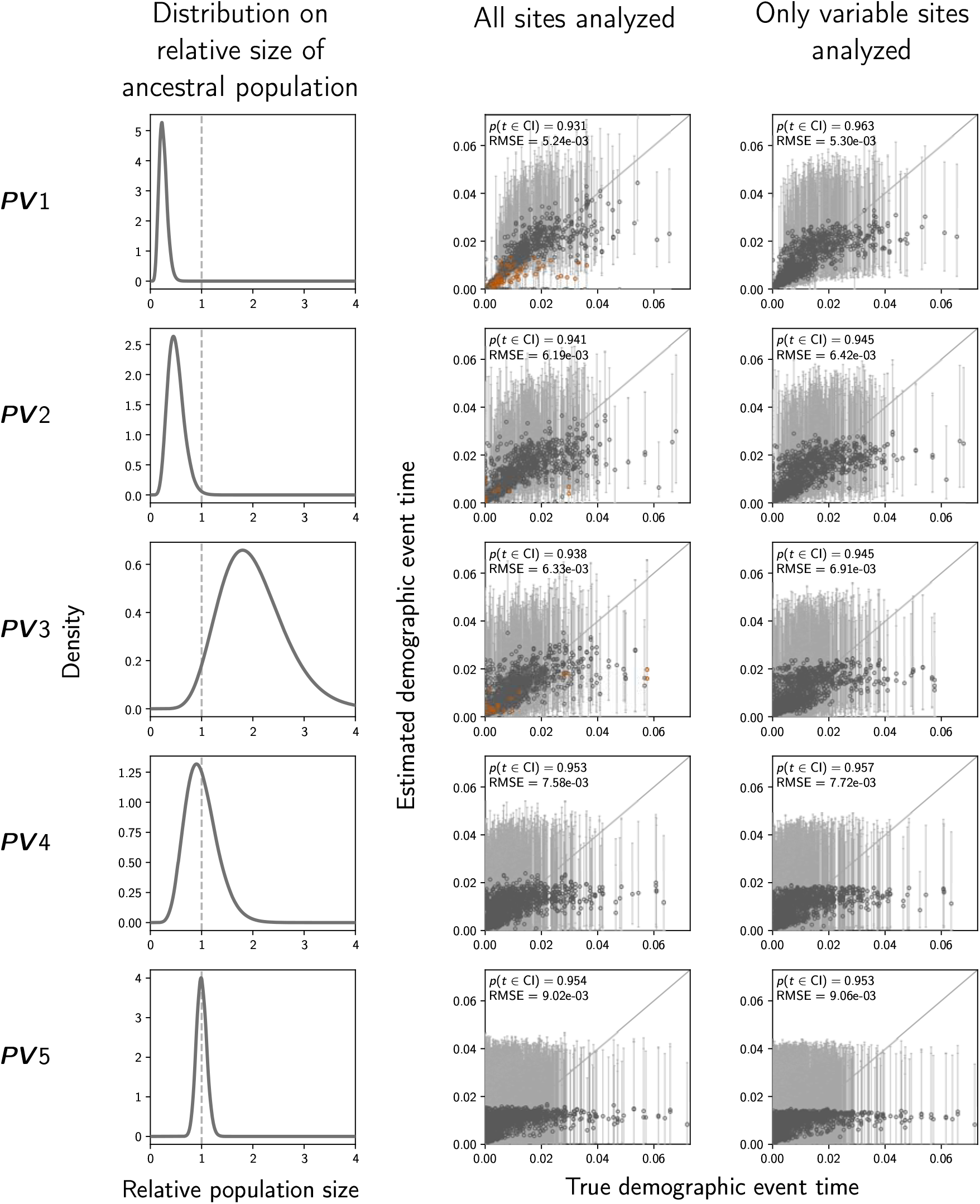
The accuracy and precision of time estimates of demographic changes (in units of expected subsitutions per site) when data were simulated and analyzed under the same distributions we initially explored (Table S1). The left column of plots shows the gamma distribution from which the relative size of the ancestral population was drawn; this was also used as the prior when each simulated data set was analyzed. The center and right column of plots show true versus estimated values when using all characters (center) or only variable characters (right). Each plotted circle and associated error bars represent the posterior mean and 95% credible interval. Estimates for which the potential-scale reduction factor was greater than 1.2 (Brooks and Gelman, 1998) are highlighted in orange. Each plot consists of 1500 estimates—500 simulated data sets, each with three demographic comparisons. For each plot, the root-mean-square error (RMSE) and the proportion of estimates for which the 95% credible interval contained the true value—*p*(*t* ∈ CI)—is given. We generated the plots using matplotlib Version 2.0.0 (Hunter, 2007).

**Figure S2.**
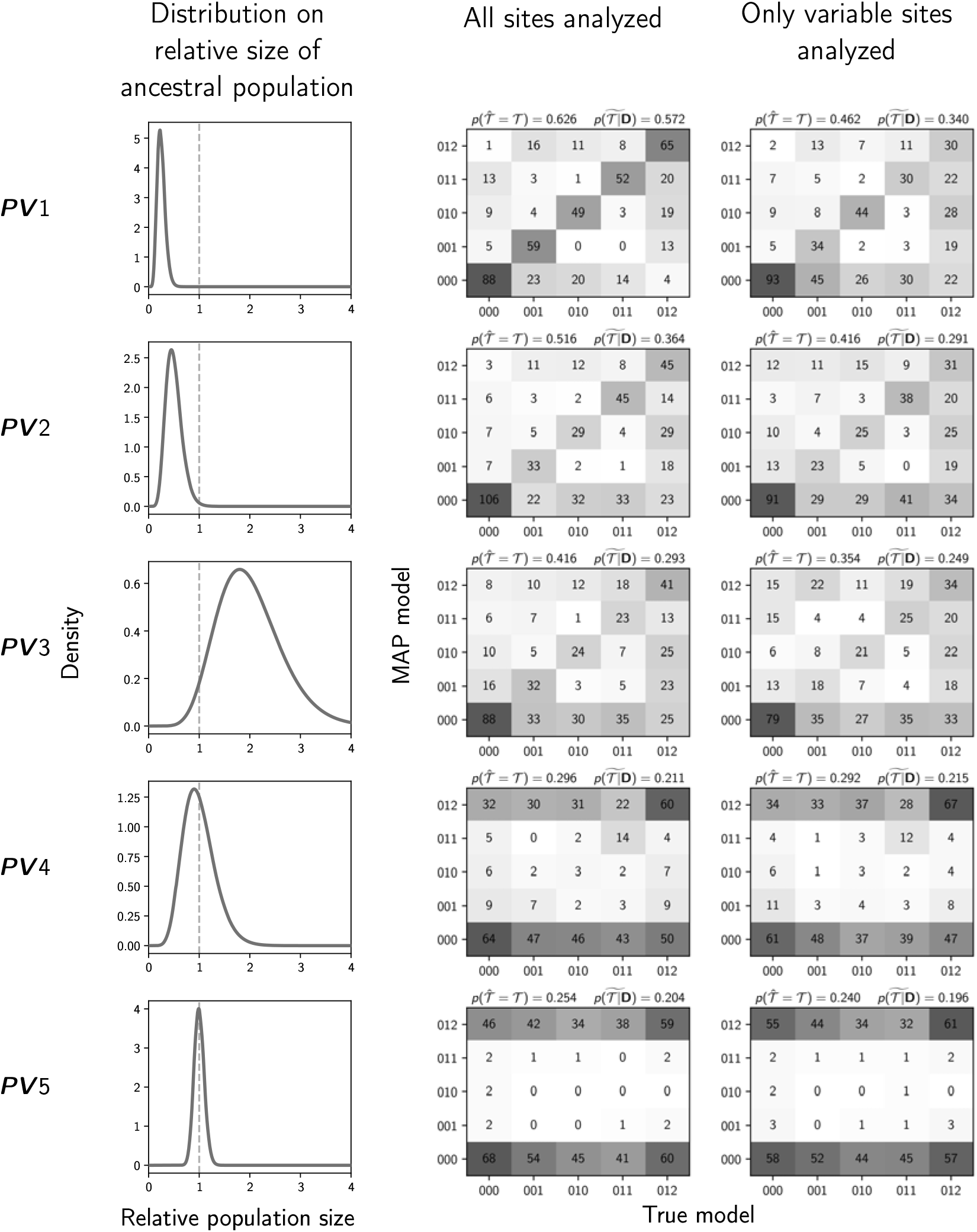
The performance of estimating the model of demographic changes when data were simulated and analyzed under the same distributions we initially explored (Table S1). The left column of plots shows the gamma distribution from which the relative size of the ancestral population was drawn; this was also used as the prior when each simulated data set was analyzed. The center and right column of plots show true versus estimated models when using all characters (center) or only variable characters (right). Each plot shows the results of the analyses of 500 simulated data sets, each with three demographic comparisons; the number of data sets that fall within each possible cell of true versus estimated model is shown, and cells with more data sets are shaded darker. Each model is represented along the plot axes by three integers that indicate the event category of each comparison (e.g., 011 represents the model in which the second and third comparison share the same event time that is distinct from the first). The estimates are based on the model with the maximum *a posteriori* probability (MAP). For each plot, the proportion of data sets for which the MAP model matched the true model—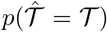—is shown in the upper left corner, and the median posterior probability of the correct model across all data sets—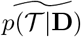—is shown in the upper right corner. We generated the plots using matplotlib Version 2.0.0 (Hunter, 2007).

**Figure S3.**
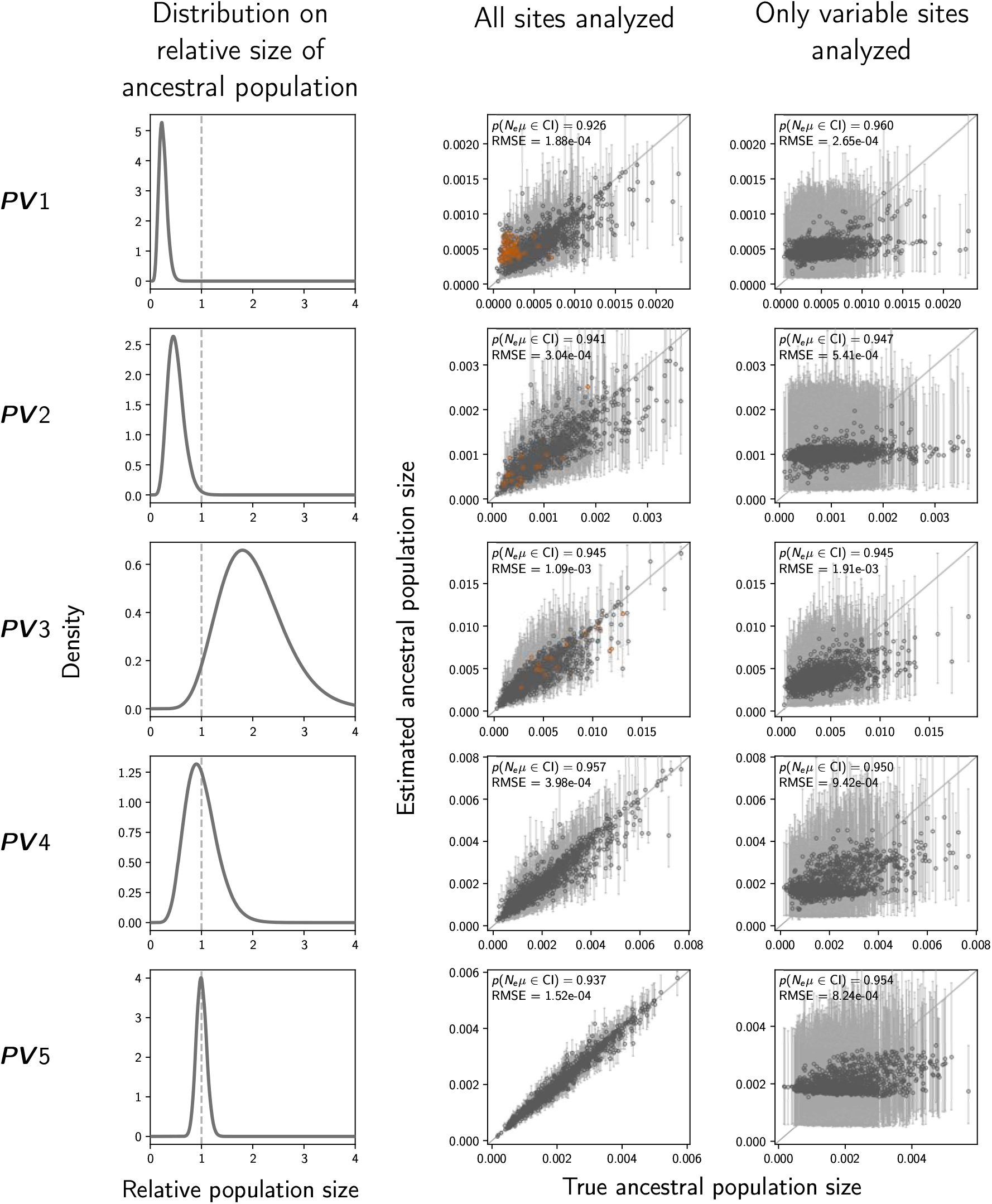
The accuracy and precision of estimates of the effective size (scaled by the mutation rate) of the population before a demographic change (“ancestral” population) when data were simulated and analyzed under the same distributions we initially explored (Table S1). The left column of plots shows the gamma distribution from which the relative size of the ancestral population was drawn; this was also used as the prior when each simulated data set was analyzed. The center and right column of plots show true versus estimated values when using all characters (center) or only variable characters (right). Each plotted circle and associated error bars represent the posterior mean and 95% credible interval. Estimates for which the potential-scale reduction factor was greater than 1.2 (Brooks and Gelman, 1998) are highlighted in orange. Each plot consists of 1500 estimates—500 simulated data sets, each with three demographic comparisons. For each plot, the root-mean-square error (RMSE) and the proportion of estimates for which the 95% credible interval contained the true value—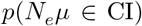—is given. We generated the plots using matplotlib Version 2.0.0 (Hunter, 2007).

**Figure S4.**
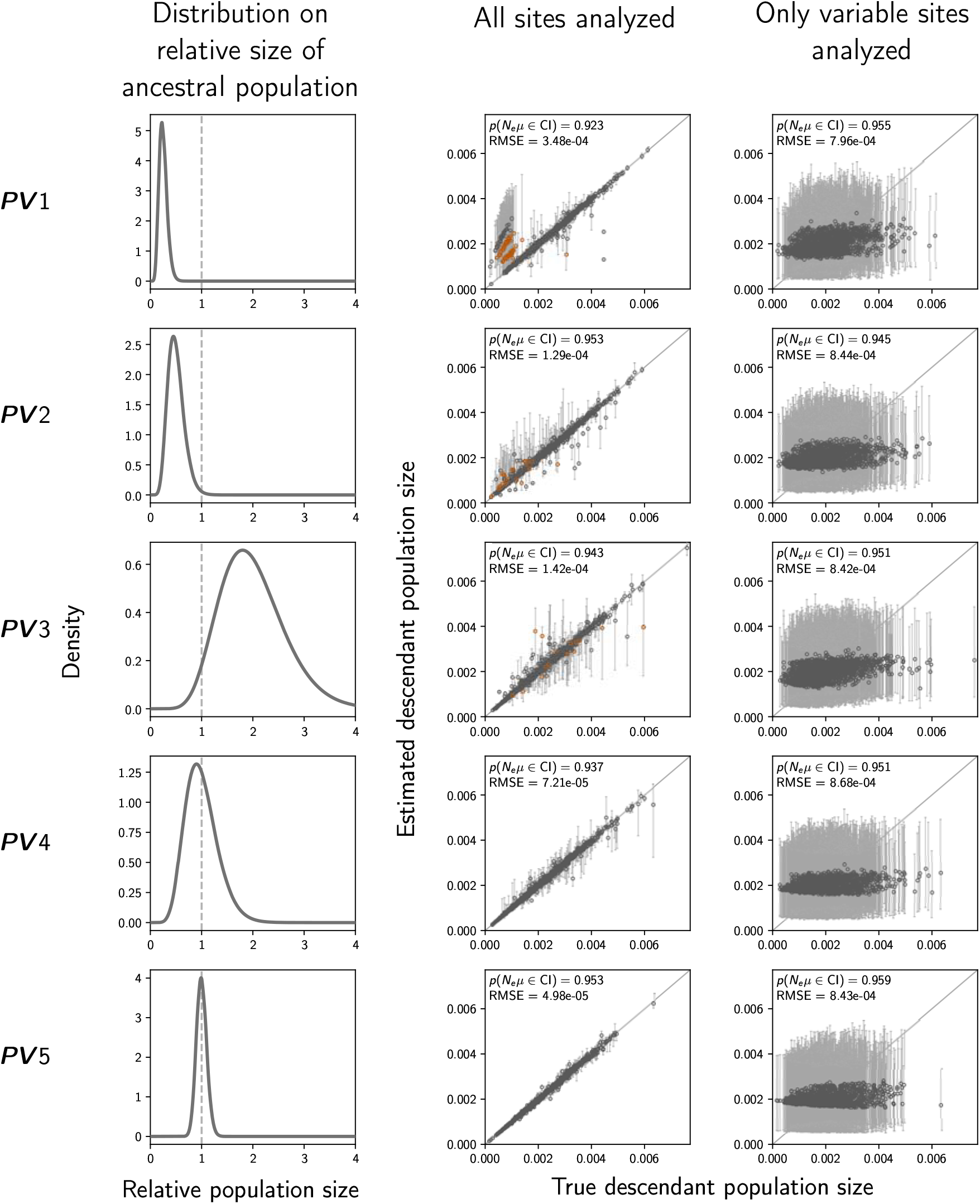
The accuracy and precision of estimates of the effective size (scaled by the mutation rate) of the population after a demographic change (“descendant” population) when data were simulated and analyzed under the same distributions we initially explored (Table S1). The left column of plots shows the gamma distribution from which the relative size of the ancestral population was drawn; this was also used as the prior when each simulated data set was analyzed. The center and right column of plots show true versus estimated values when using all characters (center) or only variable characters (right). Each plotted circle and associated error bars represent the posterior mean and 95% credible interval. Estimates for which the potential-scale reduction factor was greater than 1.2 (Brooks and Gelman, 1998) are highlighted in orange. Each plot consists of 1500 estimates—500 simulated data sets, each with three demographic comparisons. For each plot, the root-mean-square error (RMSE) and the proportion of estimates for which the 95% credible interval contained the true value—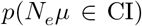—is given. We generated the plots using matplotlib Version 2.0.0 (Hunter, 2007).

**Figure S5.**
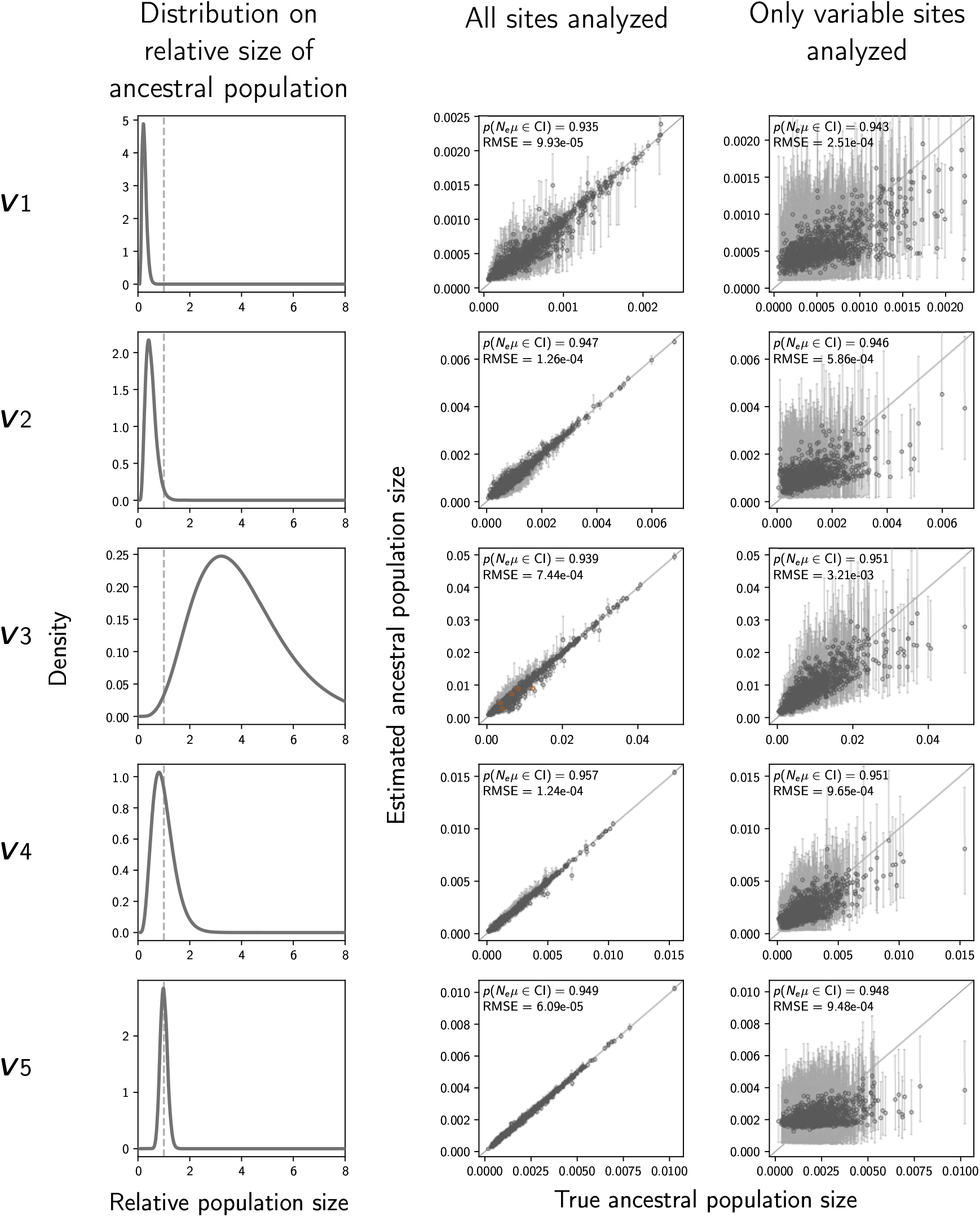
The accuracy and precision of estimates of the effective size (scaled by the mutation rate) of the population before a demographic change (“ancestral” population) when data were simulated and analyzed under the same distributions (Table 2). The left column of plots shows the gamma distribution from which the relative size of the ancestral population was drawn; this was also used as the prior when each simulated data set was analyzed. The center and right column of plots show true versus estimated values when using all characters (center) or only variable characters (right). Each plotted circle and associated error bars represent the posterior mean and 95% credible interval. Estimates for which the potential-scale reduction factor was greater than 1.2 (Brooks and Gelman, 1998) are highlighted in orange. Each plot consists of 1500 estimates—500 simulated data sets, each with three demographic comparisons. For each plot, the root-mean-square error (RMSE) and the proportion of estimates for which the 95% credible interval contained the true value—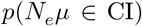—is given. We generated the plots using matplotlib Version 2.0.0 (Hunter, 2007).

**Figure S6.**
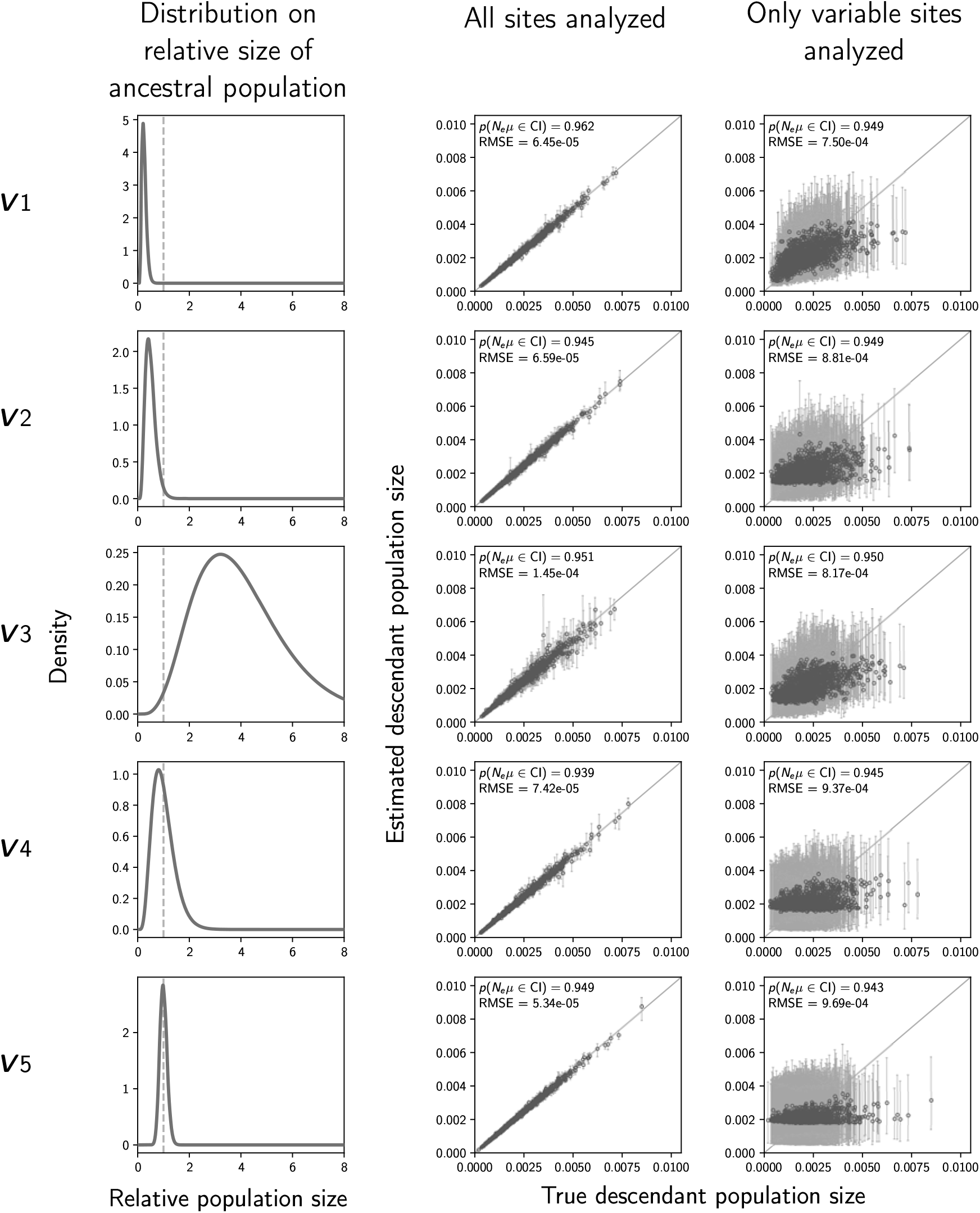
The accuracy and precision of estimates of the effective size (scaled by the mutation rate) of the population after a demographic change (“descendant” population) when data were simulated and analyzed under the same distributions (Table 2). The left column of plots shows the gamma distribution from which the relative size of the ancestral population was drawn; this was also used as the prior when each simulated data set was analyzed. The center and right column of plots show true versus estimated values when using all characters (center) or only variable characters (right). Each plotted circle and associated error bars represent the posterior mean and 95% credible interval. Estimates for which the potential-scale reduction factor was greater than 1.2 (Brooks and Gelman, 1998) are highlighted in orange. Each plot consists of 1500 estimates—500 simulated data sets, each with three demographic comparisons. For each plot, the root-mean-square error (RMSE) and the proportion of estimates for which the 95% credible interval contained the true value—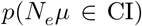—is given. We generated the plots using matplotlib Version 2.0.0 (Hunter, 2007).

**Figure S7.**
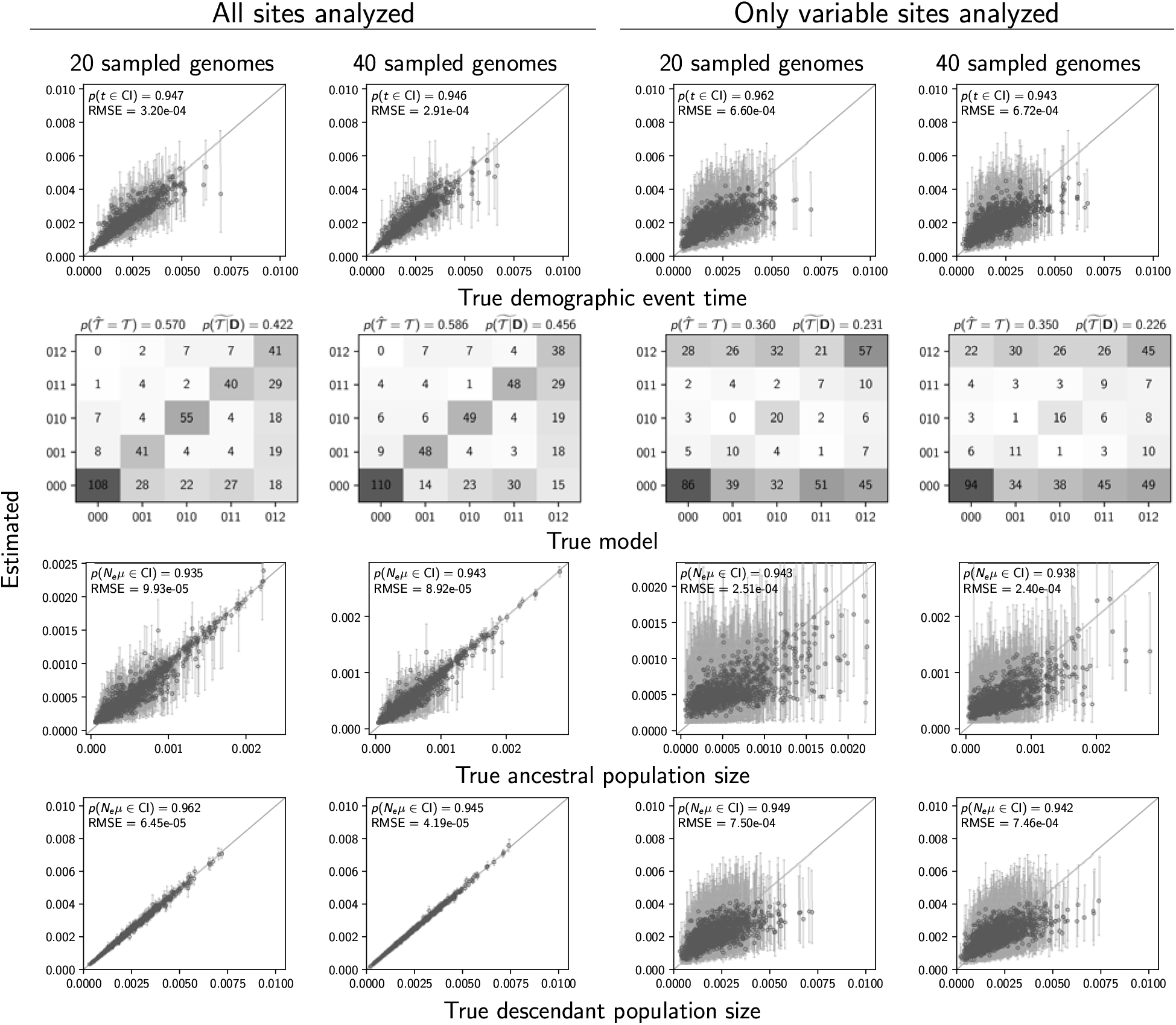
Estimates of the timing (Row 1), and sharing (Row 2) of demographic events, ancestral population size (Row 4), and descendant population size (Row 5) when 20 (Columns 1 and 3) versus 40 genomes (Columns 2 and 4) are sampled from each population. Each column plots the results from 500 data sets simulated under Condition ***V***1 (Table 2). Estimates for which the potential-scale reduction factor was greater than 1.2 (Brooks and Gelman, 1998) are highlighted in orange. We generated the plots using matplotlib Version 2.0.0 (Hunter, 2007).

**Figure S8.**
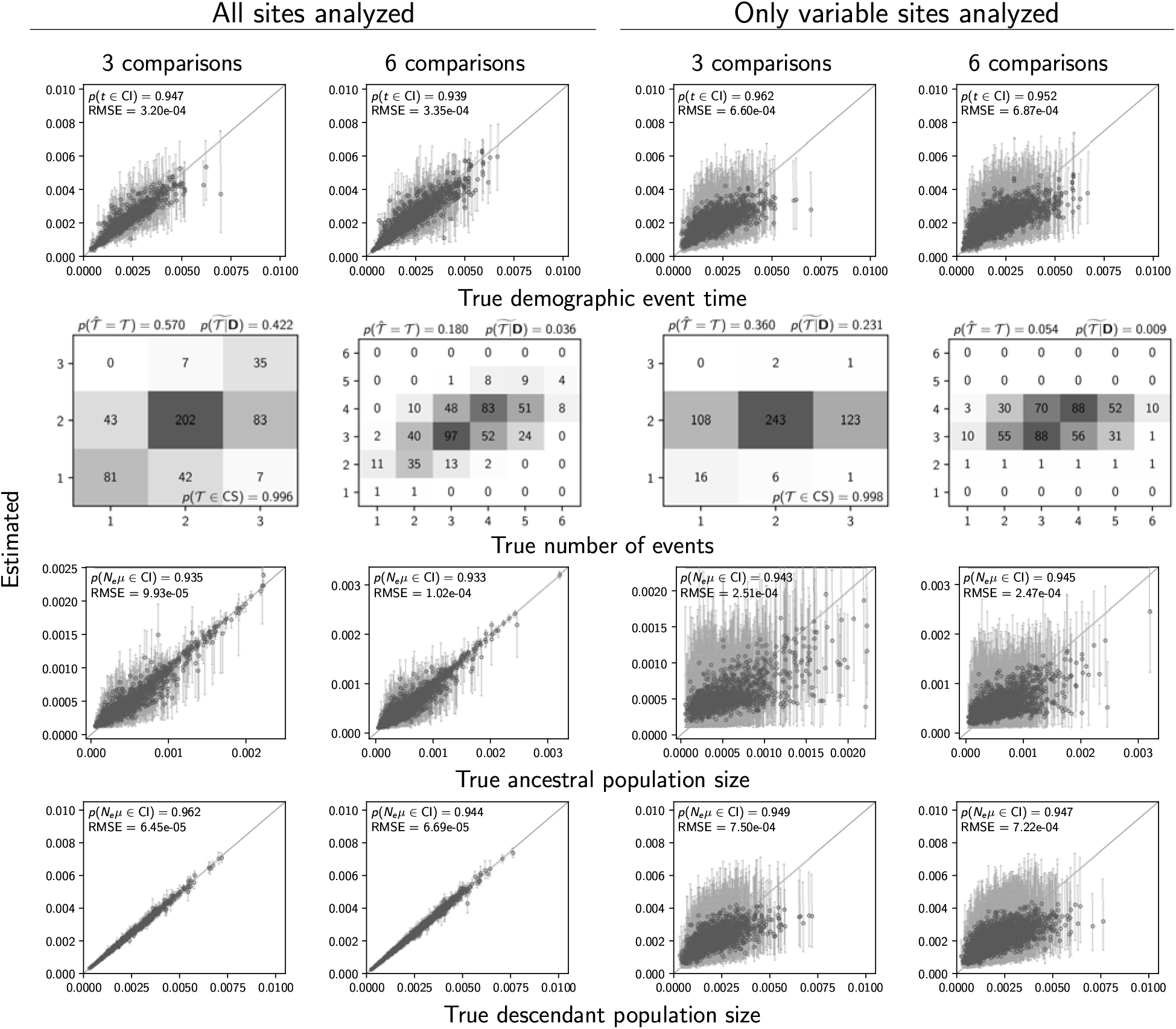
Estimates of the timing (Row 1), and sharing (Row 2) of demographic events, ancestral population size (Row 4), and descendant population size (Row 5) when three (Columns 1 and 3) versus six (Columns 2 and 4) demographic comparisons are analyzed. Each column plots the results from 500 data sets simulated under Condition ***V***1 (Table 2). Estimates for which the potential-scale reduction factor was greater than 1.2 (Brooks and Gelman, 1998) are highlighted in orange. We generated the plots using matplotlib Version 2.0.0 (Hunter, 2007).

**Figure S9.**
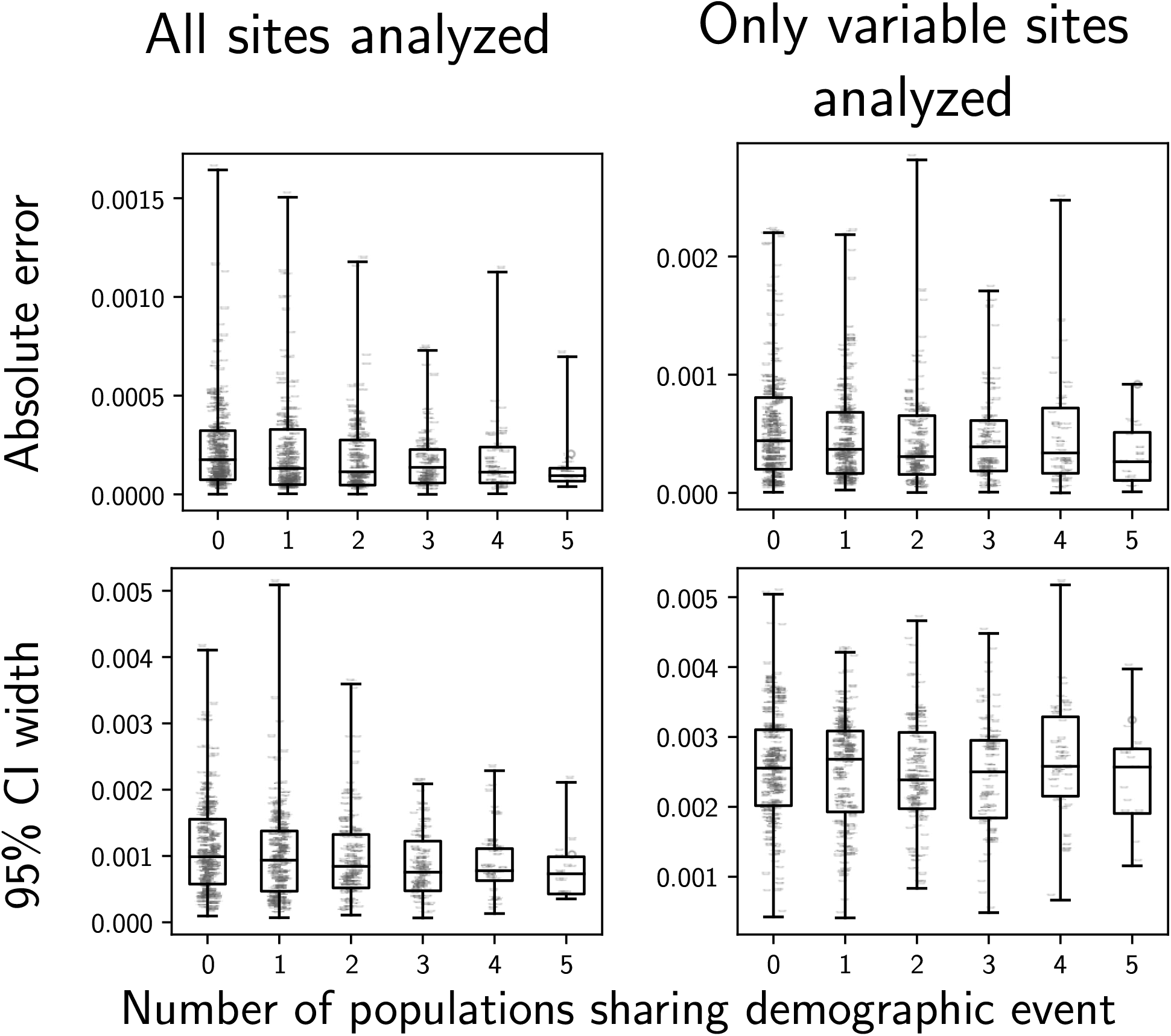
Sharing a demographic change event with more populations does not increase accuracy (as measured by absolute error; Row 1) or precision (as measured by the width of the 95% credible interval; Row 2), regardless of whether all sites (Column 1) or only variable sites (Column 2) are analyzed. Each plot shows the results from 500 data sets simulated under Condition ***V***1 (Table 2) with six demographic comparisons. We generated the plots using matplotlib Version 2.0.0 (Hunter, 2007).

**Figure S10.**
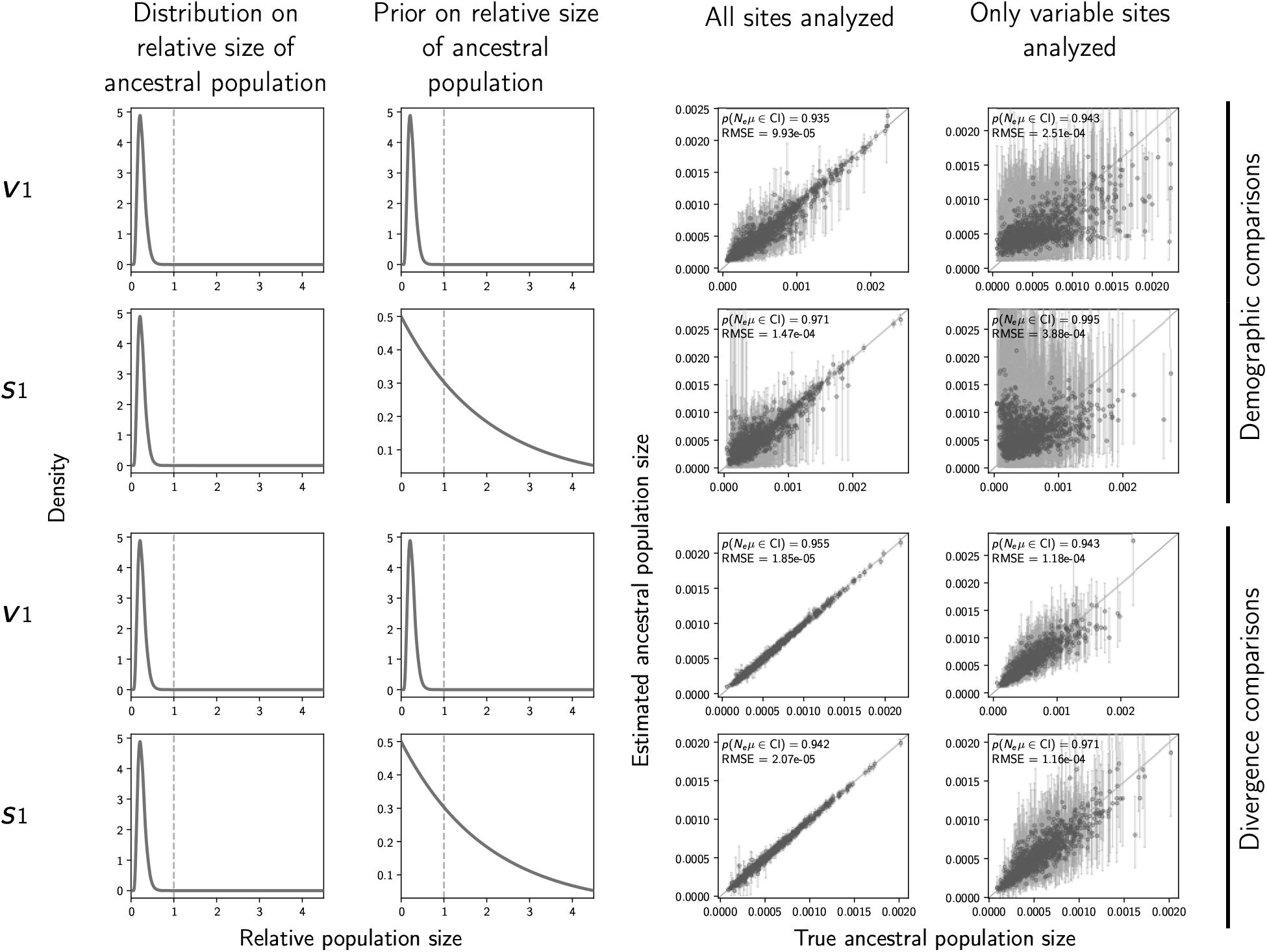
The accuracy and precision of estimates of the effective size (scaled by the mutation rate) of the ancestral population of demographic comparisons (top two rows) versus divergence comparisons (bottom two rows) when the priors are correct (first and third rows) versus when the priors are diffuse (second and fourth rows). The first and second columns of plots show the distribution on the relative effective size of the ancestral population for simulating the data (Column 1) and for the prior when analyzing the simulated data (Column 2). The third and fourth columns of plots show true versus estimated values when using all characters (Column 3) or only variable characters (Column 4). Each plotted circle and associated error bars represent the posterior mean and 95% credible interval. Estimates for which the potential-scale reduction factor was greater than 1.2 (Brooks and Gelman, 1998) are highlighted in orange. Each plot comprises 500 simulated data sets, each with three demographic comparisons (Rows 1–2) or divergence comparisons (Rows 3–4). For each plot, the root-mean-square error (RMSE) and the proportion of estimates for which the 95% credible interval contained the true value—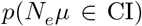—is given. The first row of plots are repeated from Figure S5 for comparison. We generated the plots using matplotlib Version 2.0.0 (Hunter, 2007).

**Figure S11.**
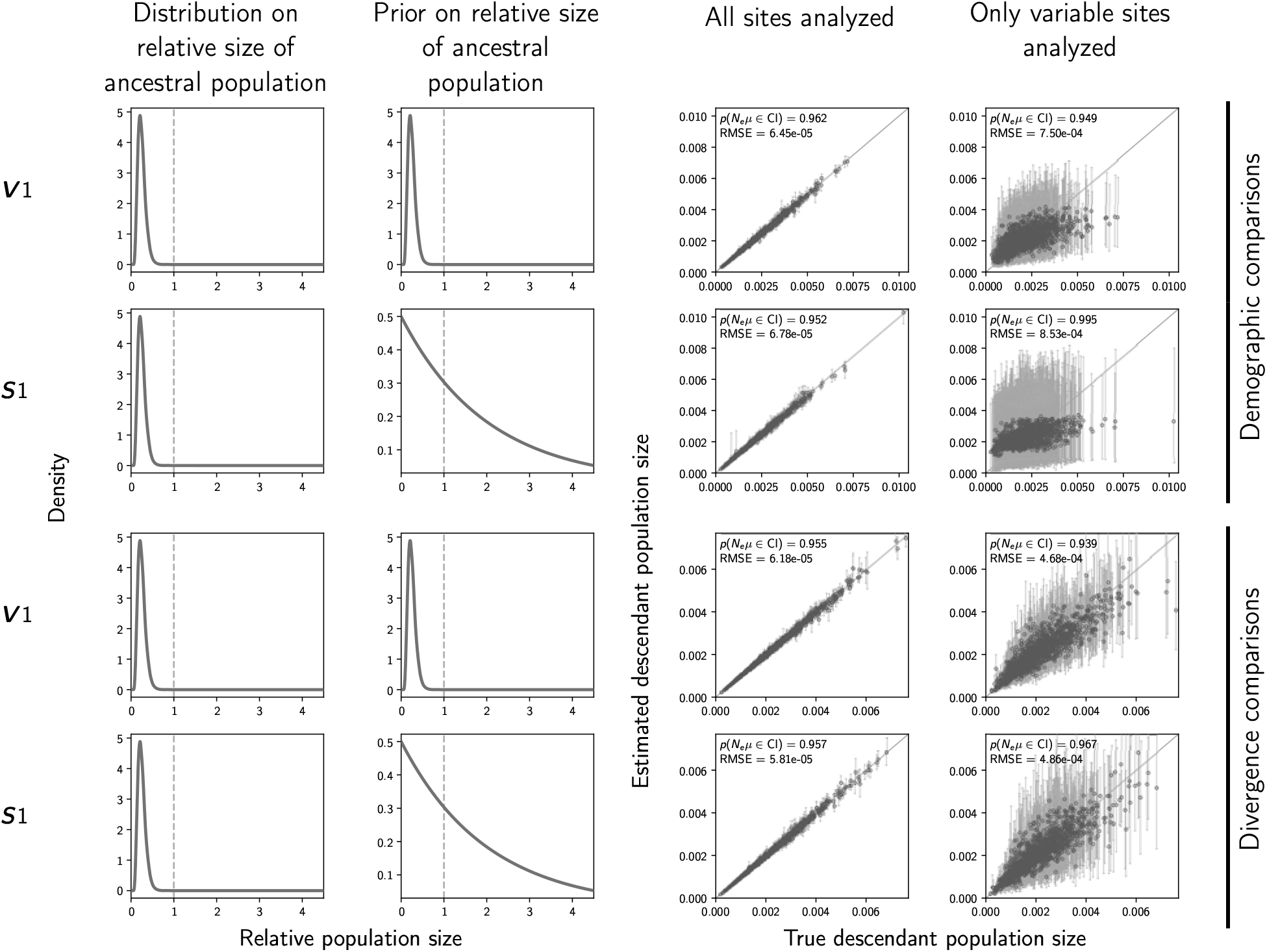
The accuracy and precision of estimates of the effective size (scaled by the mutation rate) of the descendant population(s) of demographic comparisons (top two rows) versus divergence comparisons (bottom two rows) when the priors are correct (first and third rows) versus when the priors are diffuse (second and fourth rows). The first and second columns of plots show the distribution on the relative effective size of the ancestral population for simulating the data (Column 1) and for the prior when analyzing the simulated data (Column 2). The third and fourth columns of plots show true versus estimated values when using all characters (Column 3) or only variable characters (Column 4). Each plotted circle and associated error bars represent the posterior mean and 95% credible interval. Estimates for which the potential-scale reduction factor was greater than 1.2 (Brooks and Gelman, 1998) are highlighted in orange. Each plot comprises 500 simulated data sets, each with three demographic comparisons (Rows 1–2) or divergence comparisons (Rows 3–4). For each plot, the root-mean-square error (RMSE) and the proportion of estimates for which the 95% credible interval contained the true value—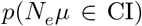—is given. The first row of plots are repeated from Figure S6 for comparison. We generated the plots using matplotlib Version 2.0.0 (Hunter, 2007).

**Figure S12.**
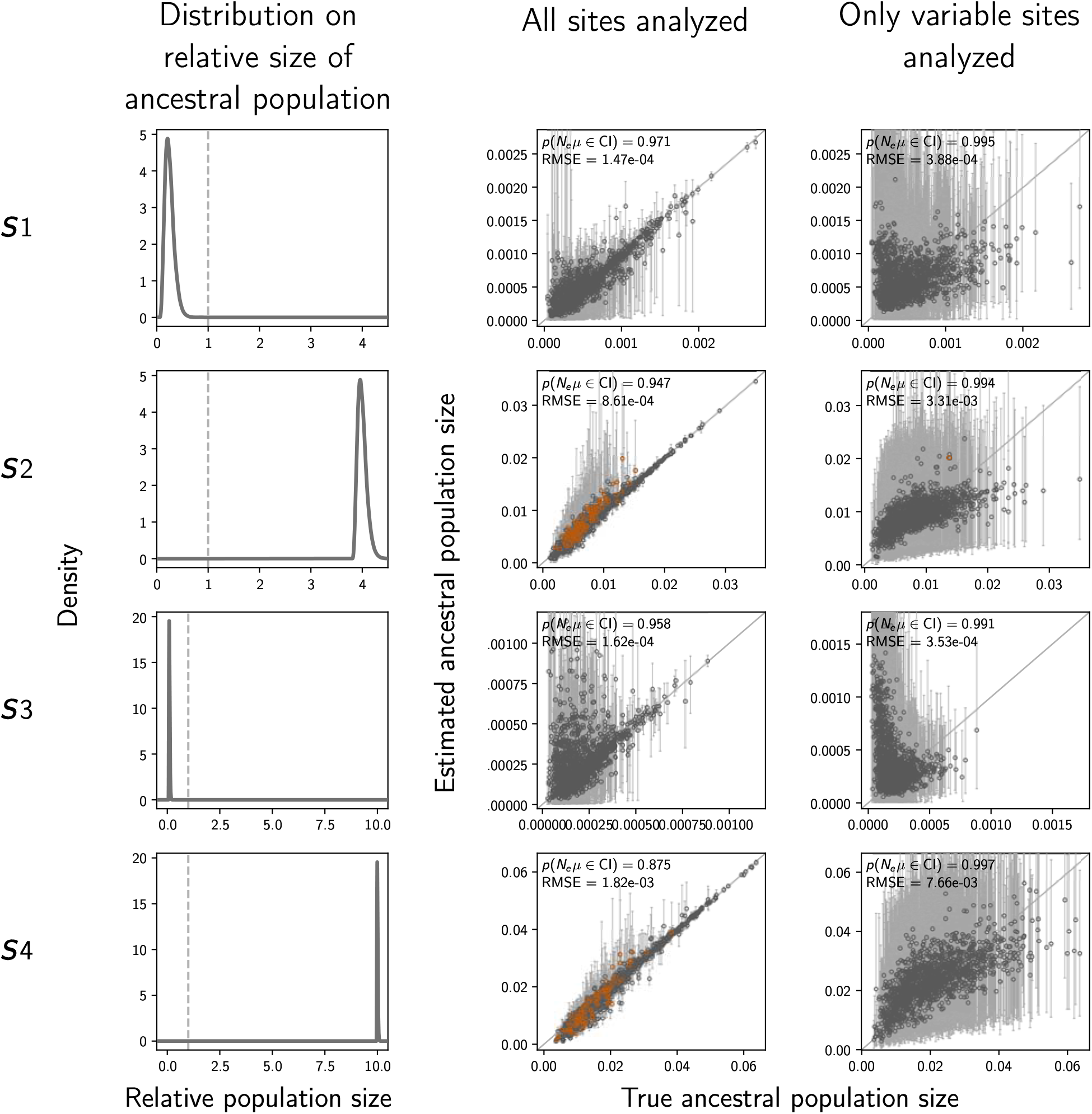
The accuracy and precision of estimates of the effective size of the population before the demographic change (i.e., ancestral population) when the prior distributions are diffuse (Conditions ***S***1–***S***4; Table 2). The first column of plots shows the distribution on the relative effective size of the ancestral population under which the data were simulated, and the second and third columns of plots show true versus estimated values when using all characters (Column 2) or only variable characters (Column 3). Each plotted circle and associated error bars represent the posterior mean and 95% credible interval. Estimates for which the potential-scale reduction factor was greater than 1.2 (Brooks and Gelman, 1998) are highlighted in orange. Each plot comprises 500 simulated data sets, each with three demographic comparisons. For each plot, the root-mean-square error (RMSE) and the proportion of estimates for which the 95% credible interval contained the true value—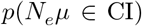—is given. The first row of plots are repeated from Figure S10 for comparison. We generated the plots using matplotlib Version 2.0.0 (Hunter, 2007).

**Figure S13.**
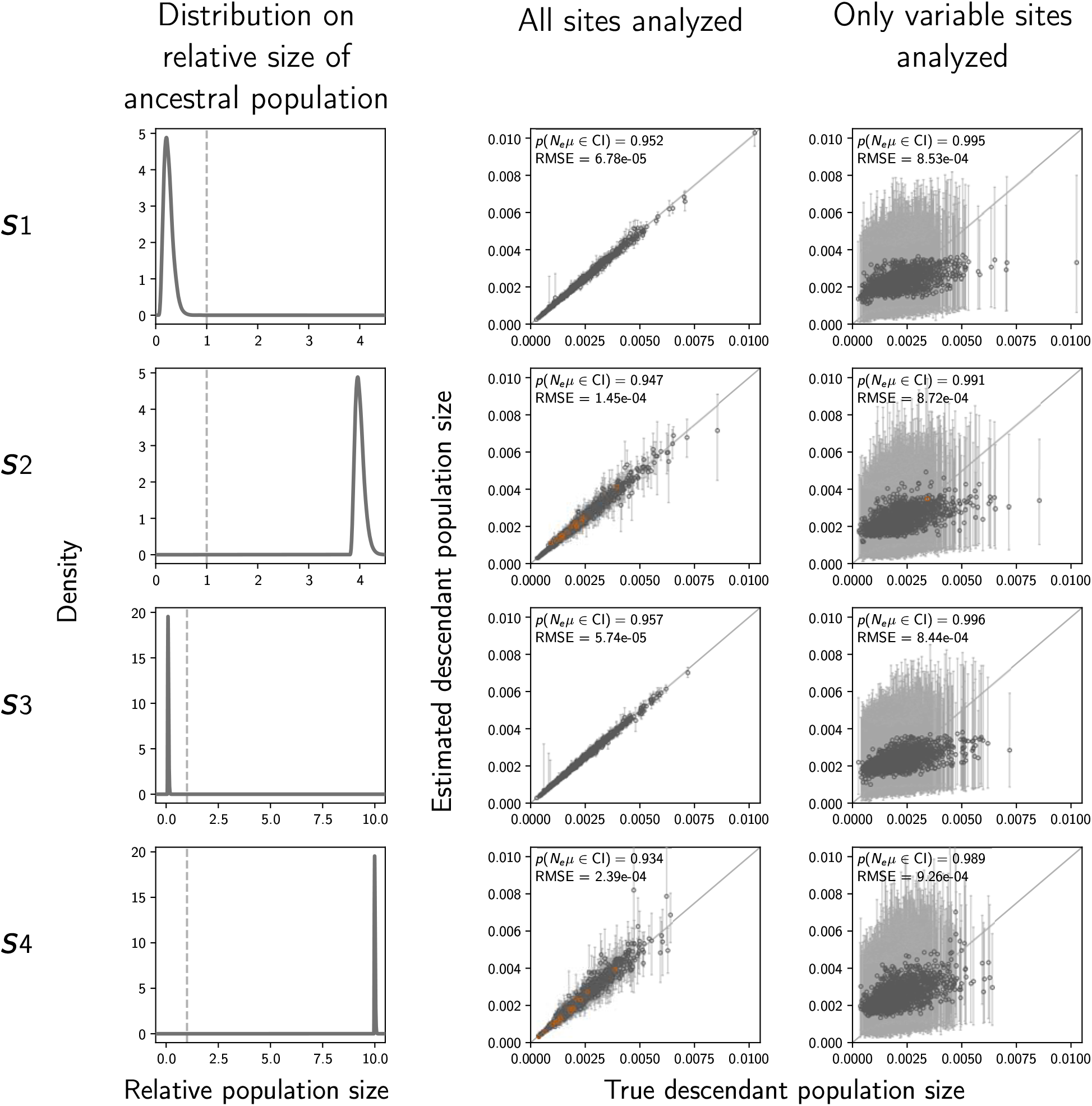
The accuracy and precision of estimates of the effective size of the population after the demographic change (i.e., descendant population) when the prior distributions are diffuse (Conditions ***S***1–***S***4; Table 2). The first column of plots shows the distribution on the relative effective size of the ancestral population under which the data were simulated, and the second and third columns of plots show true versus estimated values when using all characters (Column 2) or only variable characters (Column 3). Each plotted circle and associated error bars represent the posterior mean and 95% credible interval. Estimates for which the potential-scale reduction factor was greater than 1.2 (Brooks and Gelman, 1998) are highlighted in orange. Each plot comprises 500 simulated data sets, each with three demographic comparisons. For each plot, the root-mean-square error (RMSE) and the proportion of estimates for which the 95% credible interval contained the true value—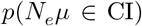—is given. The first row of plots are repeated from Figure S11 for comparison. We generated the plots using matplotlib Version 2.0.0 (Hunter, 2007).

**Figure S14.**
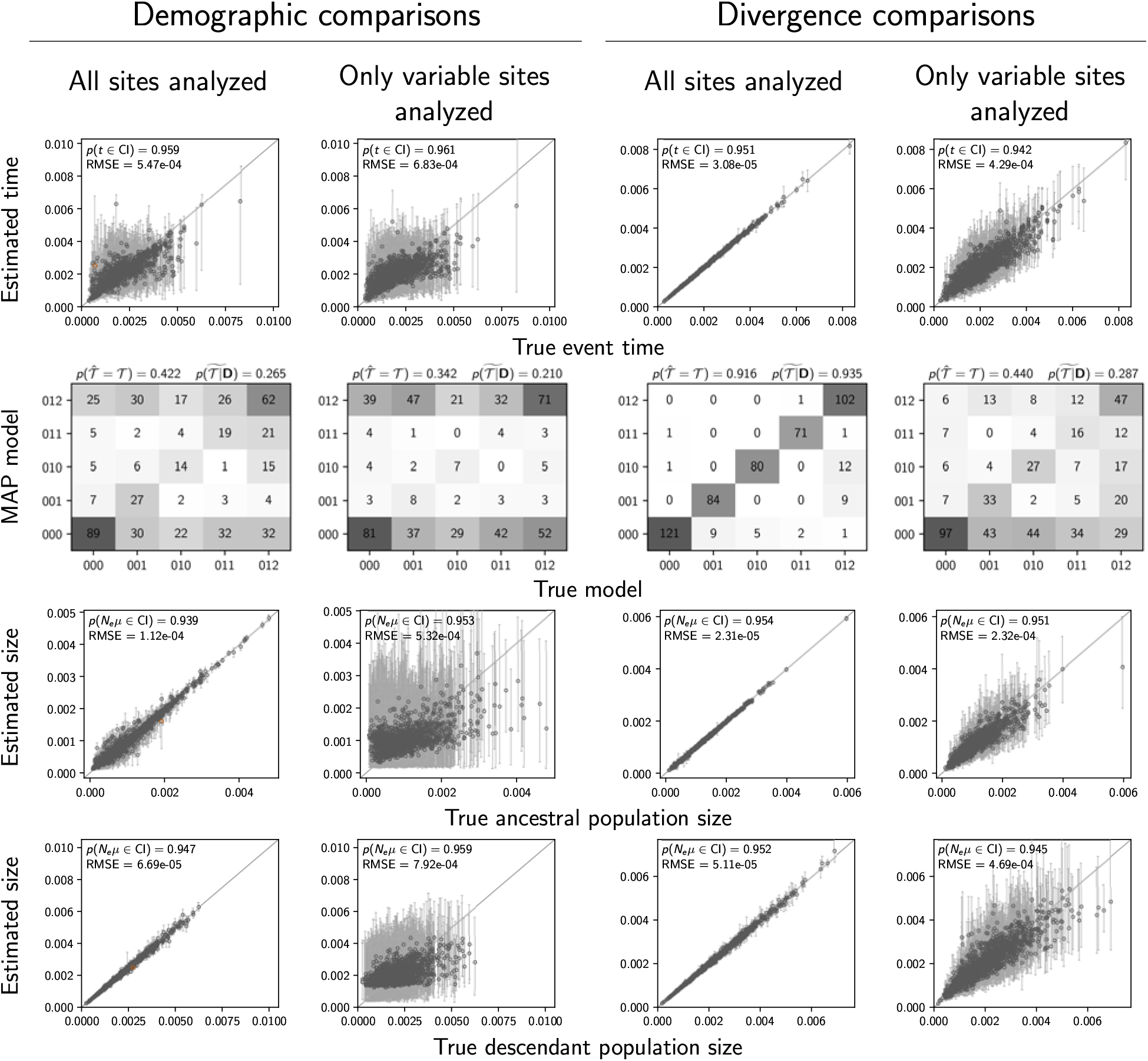
Results of analyses of 500 data sets simulated with six comparisons comprising a mix of three populations that experienced a demographic change and three pairs of populations that diverged. The performance of estimating the timing of events (Row 1), sharing of events (Rows 2–3), ancestral population size (Row 4), and descendant population size (Row 5) are shown separately for the three populations that experienced a demographic change (Columns 1 and 2) and the three pairs of populations that diverged (Columns 3 and 4). The plots of the demographic comparisons (Columns 1 and 2) are comparable to the second column of Figures 2, 3, S5, and S6; the same priors on event times and ancestral population size were used. Estimates for which the potential-scale reduction factor was greater than 1.2 (Brooks and Gelman, 1998) are highlighted in orange. We generated the plots using matplotlib Version 2.0.0 (Hunter, 2007).

**Figure S15.**
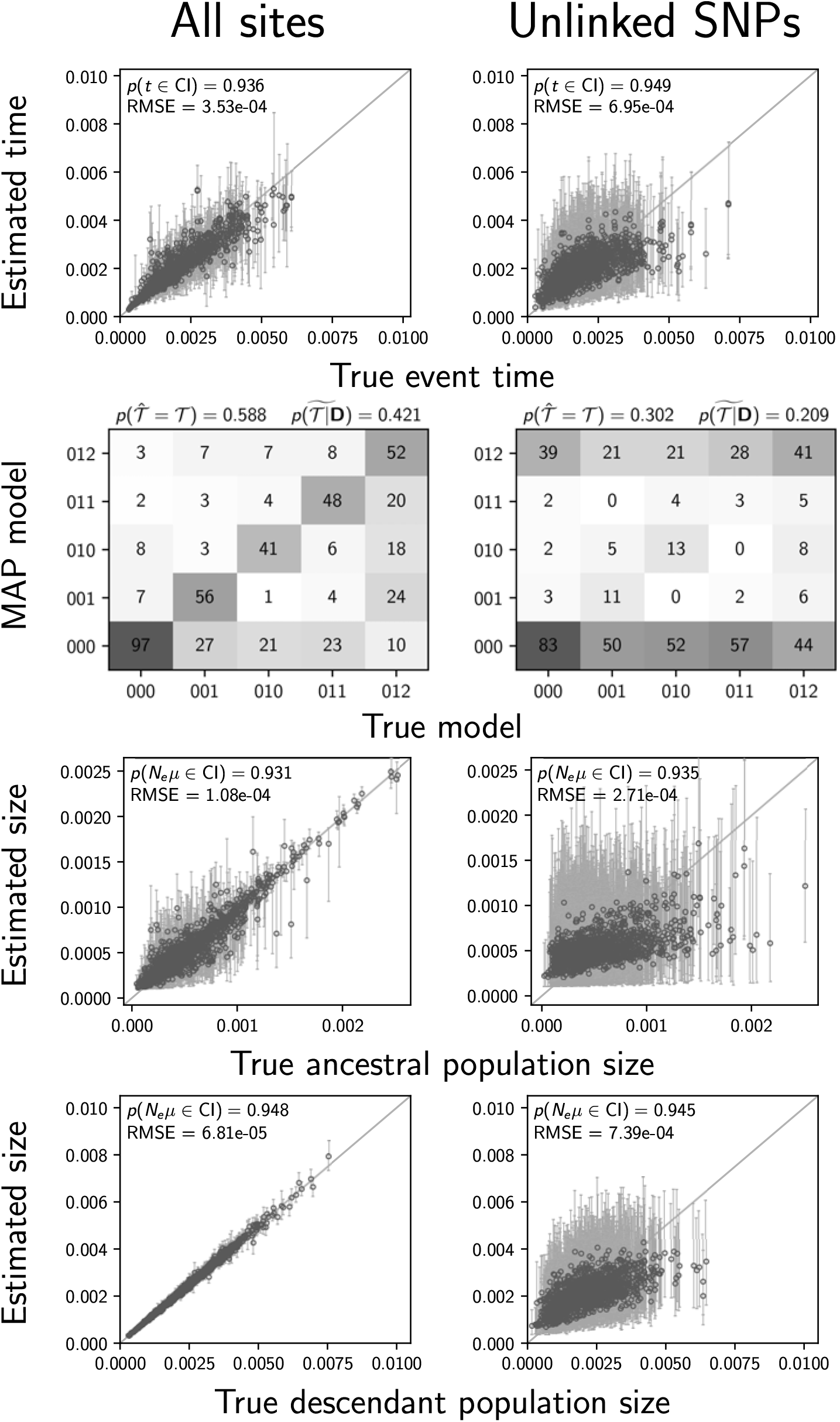
Estimates of the timing (Row 1), and sharing (Row 2) of demographic events, ancestral population size (Row 4), and descendant population size (Row 5) when using all characters (left column) or only unlinked variable characters (right column) from data sets simulated with 5000 loci of 100 linked bases from three demographic comparisons. The plots are comparable to the first row of Figures 2, 3, S5, and S6; the only difference is the linkage of characters into loci. Estimates for which the potential-scale reduction factor was greater than 1.2 (Brooks and Gelman, 1998) are highlighted in orange. Each plot shows the results from 500 simulated data sets, each with three demographic comparisons. We generated the plots using matplotlib Version 2.0.0 (Hunter, 2007)

**Figure S16.**
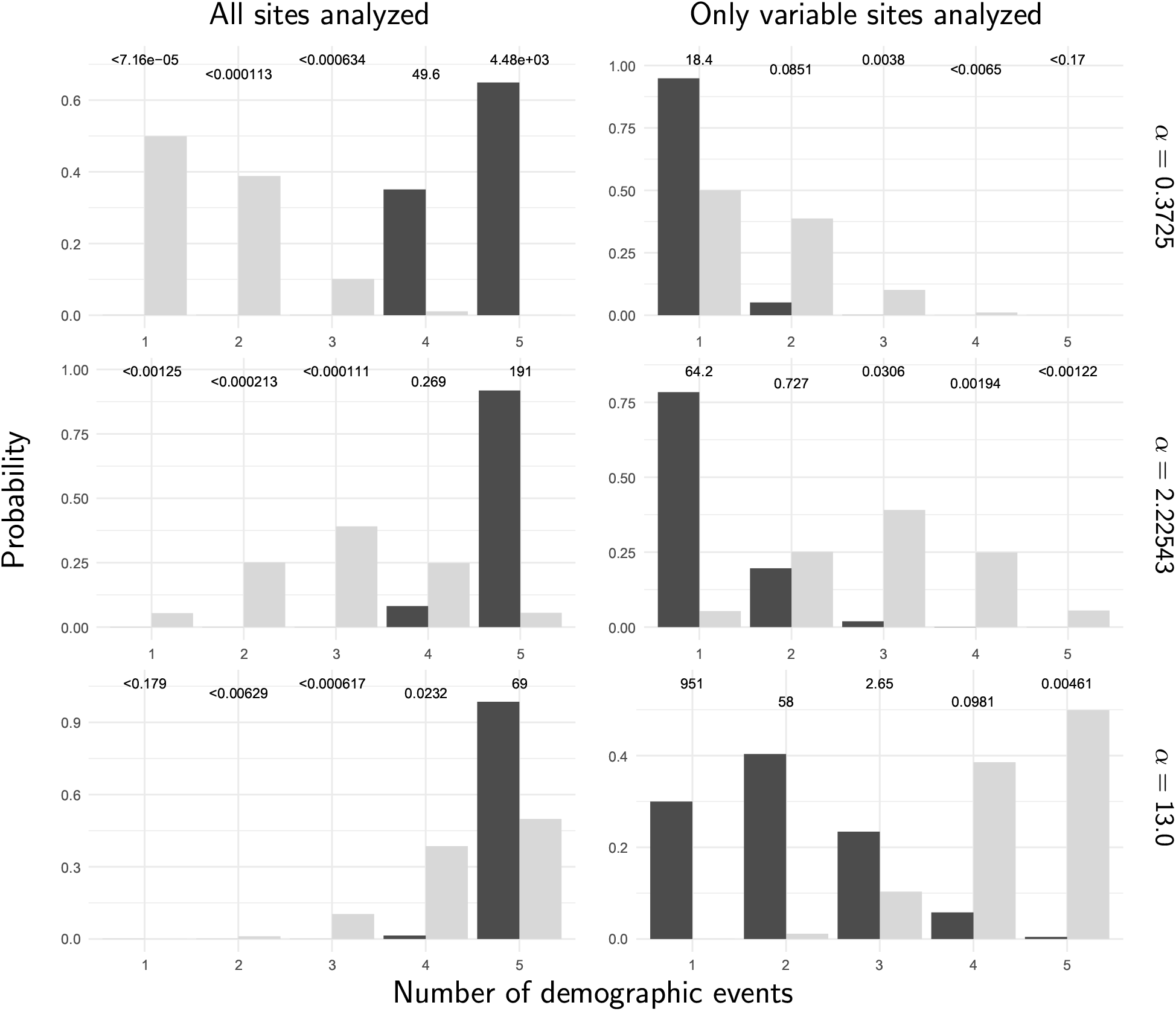
The prior (light bars) and posterior (dark bars) probabilities of the number of demographic events across five stickleback populations when all of the sites (left column) or only variable sites (right column) of the RADseq alignments are analyzed. Each row shows results under a different prior on the concentration parameter of the dirichlet process. We generated the plots with ggplot2 Version 2.2.1 (Wickham, 2009).

**Figure S17.**
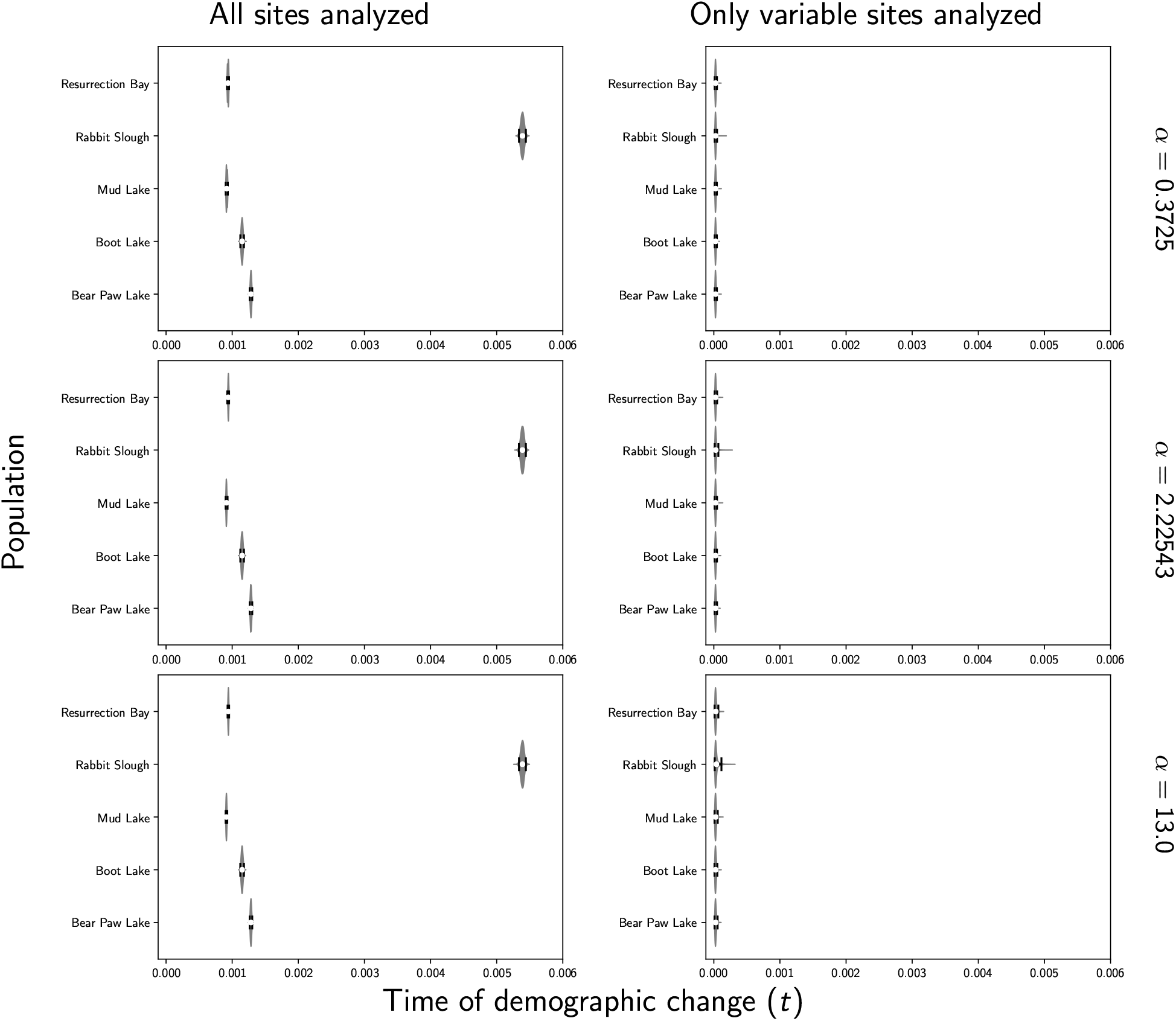
Estimates of the time of a change in population size across five stickleback populations when all of the sites (left column) or only variable sites (right column) of the RADseq alignments are analyzed. Each row shows results under a different prior on the concentration parameter of the dirichlet process. We generated the plots using matplotlib Version 2.0.0 (Hunter, 2007).

**Figure S18.**
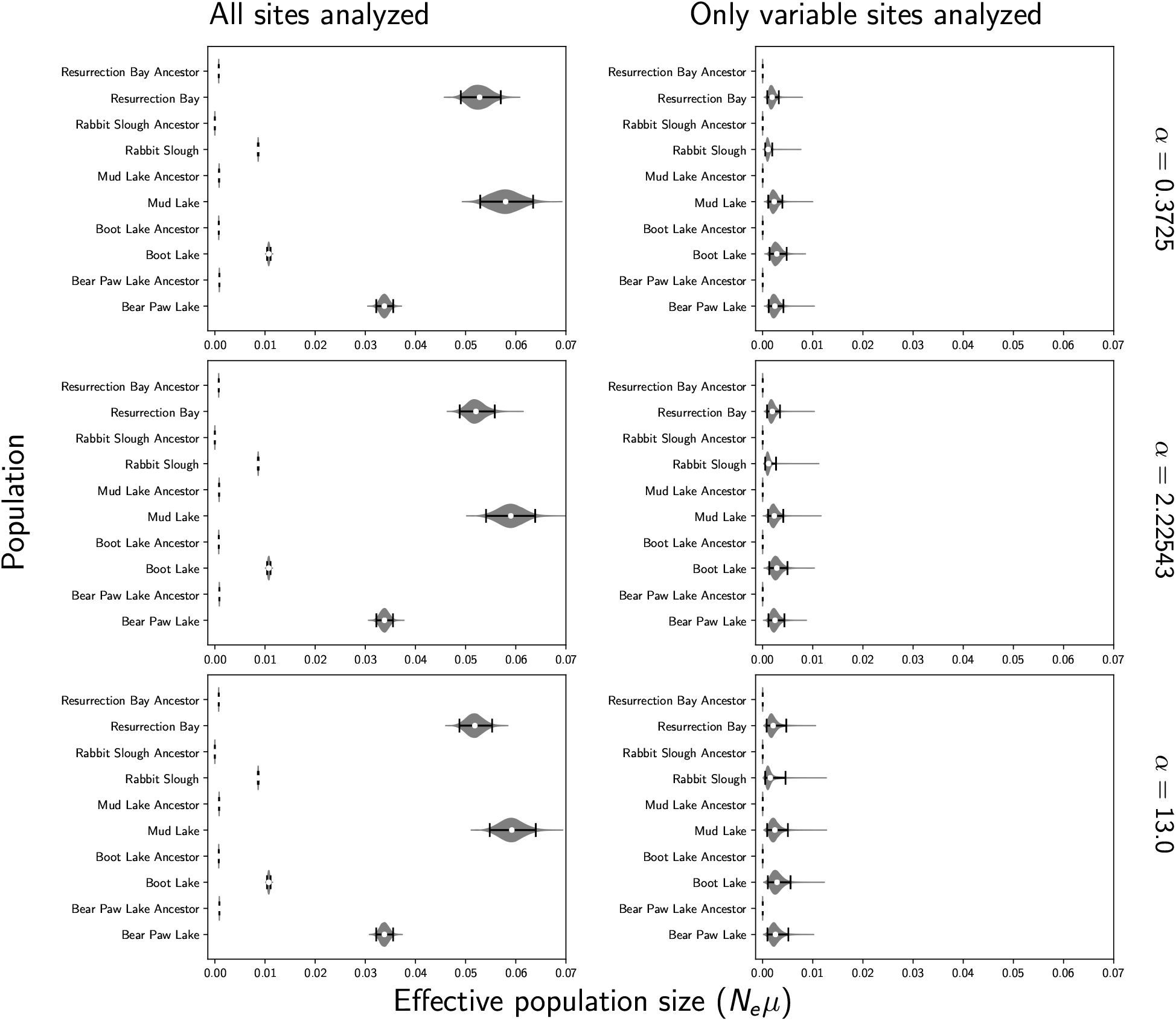
Estimates of the effective population size before (“ancestor”) and after a demographic change across five stickleback populations when all of the sites (left column) or only variable sites (right column) of the RADseq alignments are analyzed. Each row shows results under a different prior on the concentration parameter of the dirichlet process. We generated the plots using matplotlib Version 2.0.0 (Hunter, 2007).

**Figure S19.**
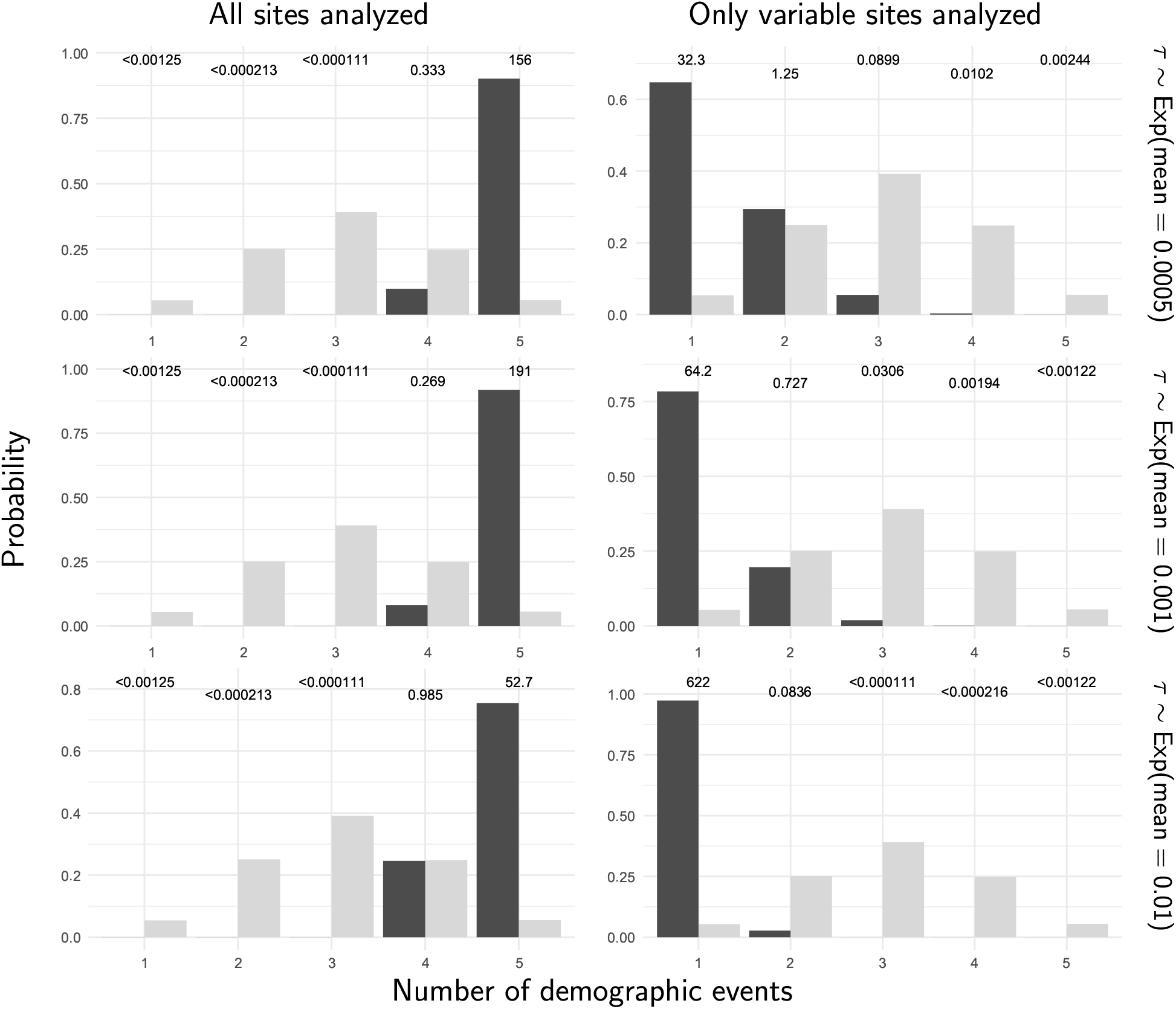
The prior (light bars) and posterior (dark bars) probabilities of the number of demographic events across five stickleback populations when all of the sites (left column) or only variable sites (right column) of the RADseq alignments are analyzed. Each row shows results under a different prior on the timing of the change in population size. We generated the plots with ggplot2 Version 2.2.1 (Wickham, 2009).

**Figure S20.**
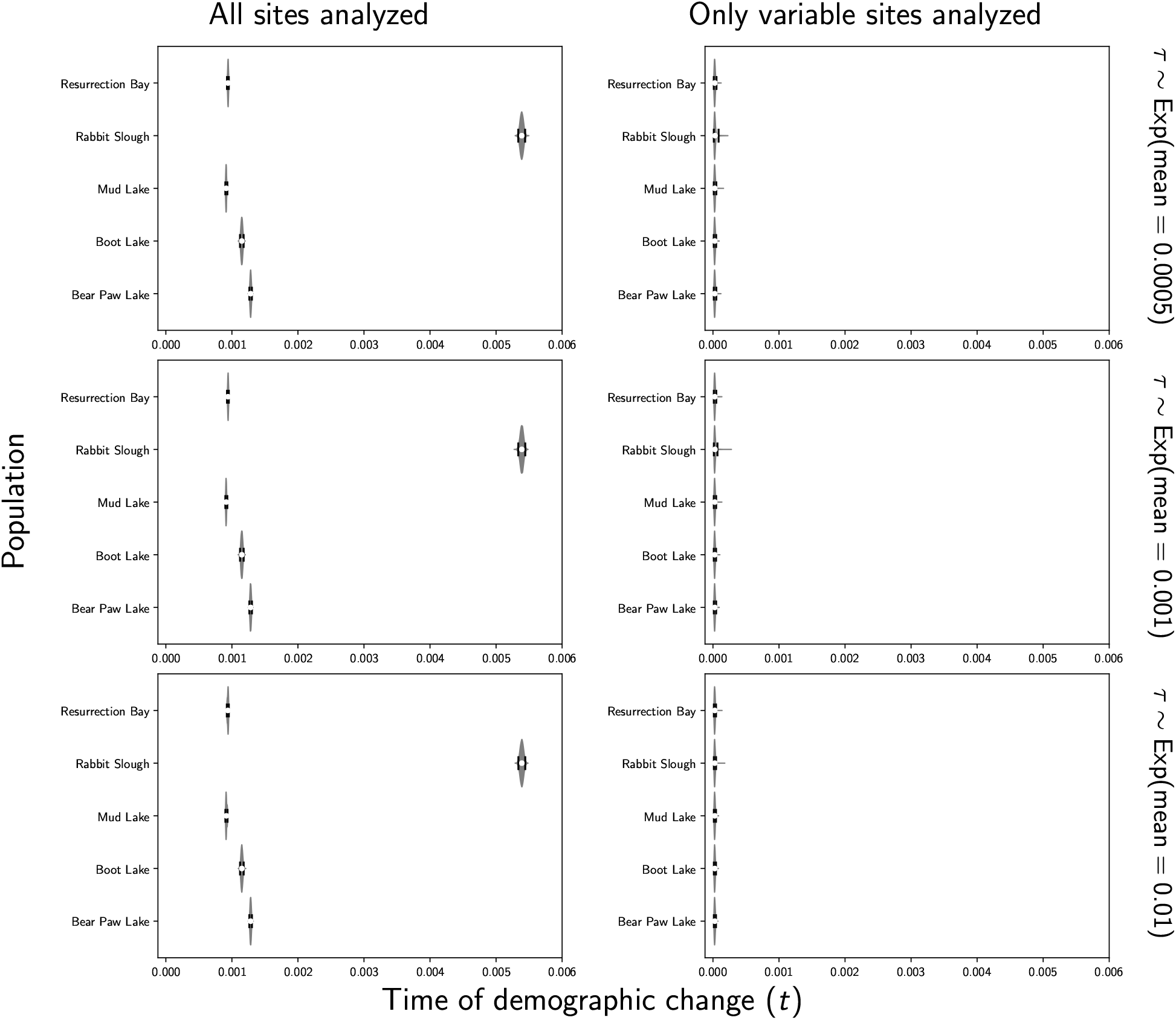
Estimates of the time of a change in population size across five stickleback populations when all of the sites (left column) or only variable sites (right column) of the RADseq alignments are analyzed. Each row shows results under a different prior on the timing of the change in population size. We generated the plots using matplotlib Version 2.0.0 (Hunter, 2007).

**Figure S21.**
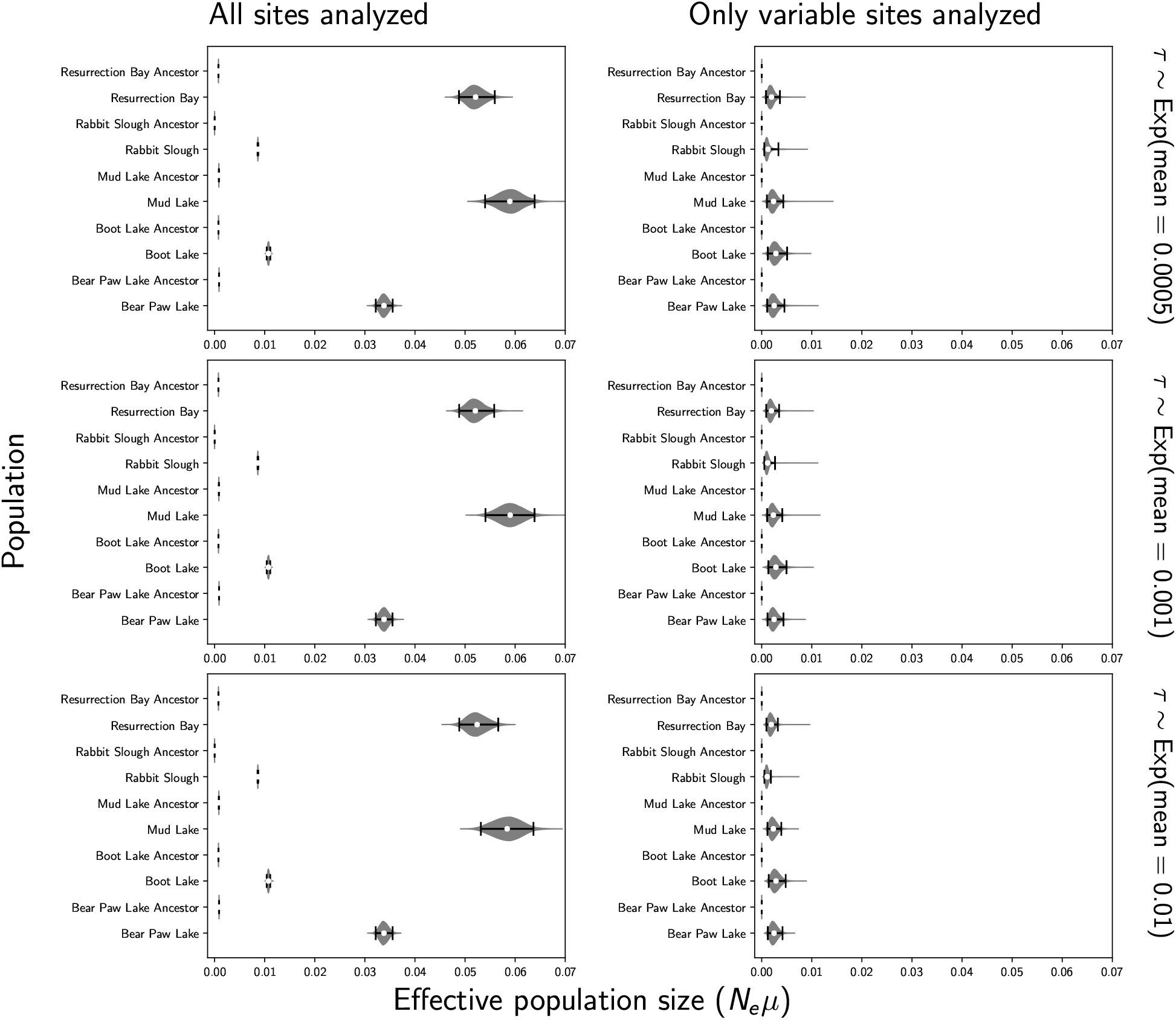
Estimates of the effective population size before (“ancestor”) and after a demographic change across five stickleback populations when all of the sites (left column) or only variable sites (right column) of the RADseq alignments are analyzed. Each row shows results under a different prior on the timing of the change in population size. We generated the plots using matplotlib Version 2.0.0 (Hunter, 2007).

**Figure S22.**
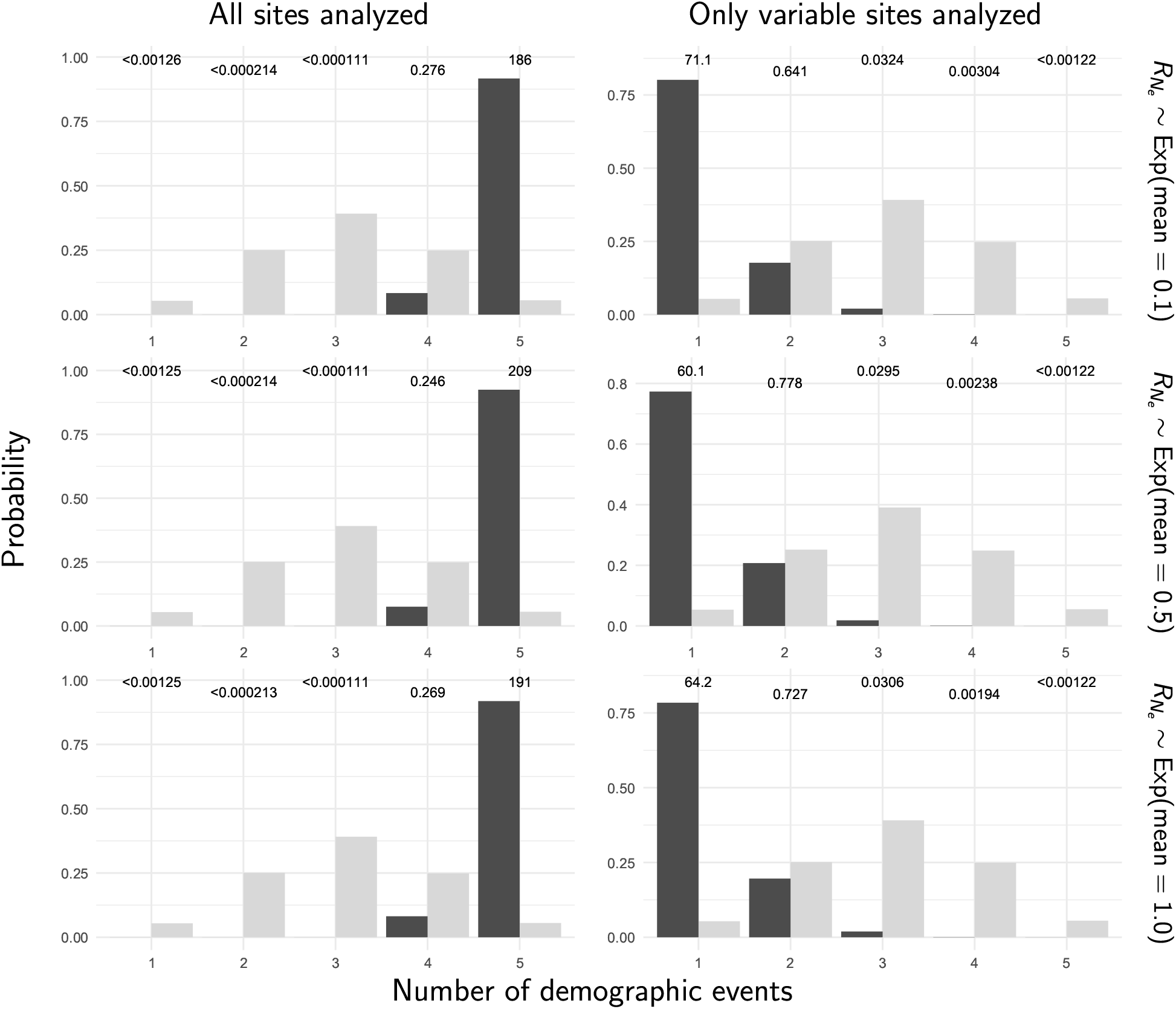
The prior (light bars) and posterior (dark bars) probabilities of the number of demographic events across five stickleback populations when all of the sites (left column) or only variable sites (right column) of the RADseq alignments are analyzed. Each row shows results under a different prior on the relative effective size of the ancestral population. We generated the plots with ggplot2 Version 2.2.1 (Wickham, 2009).

**Figure S23.**
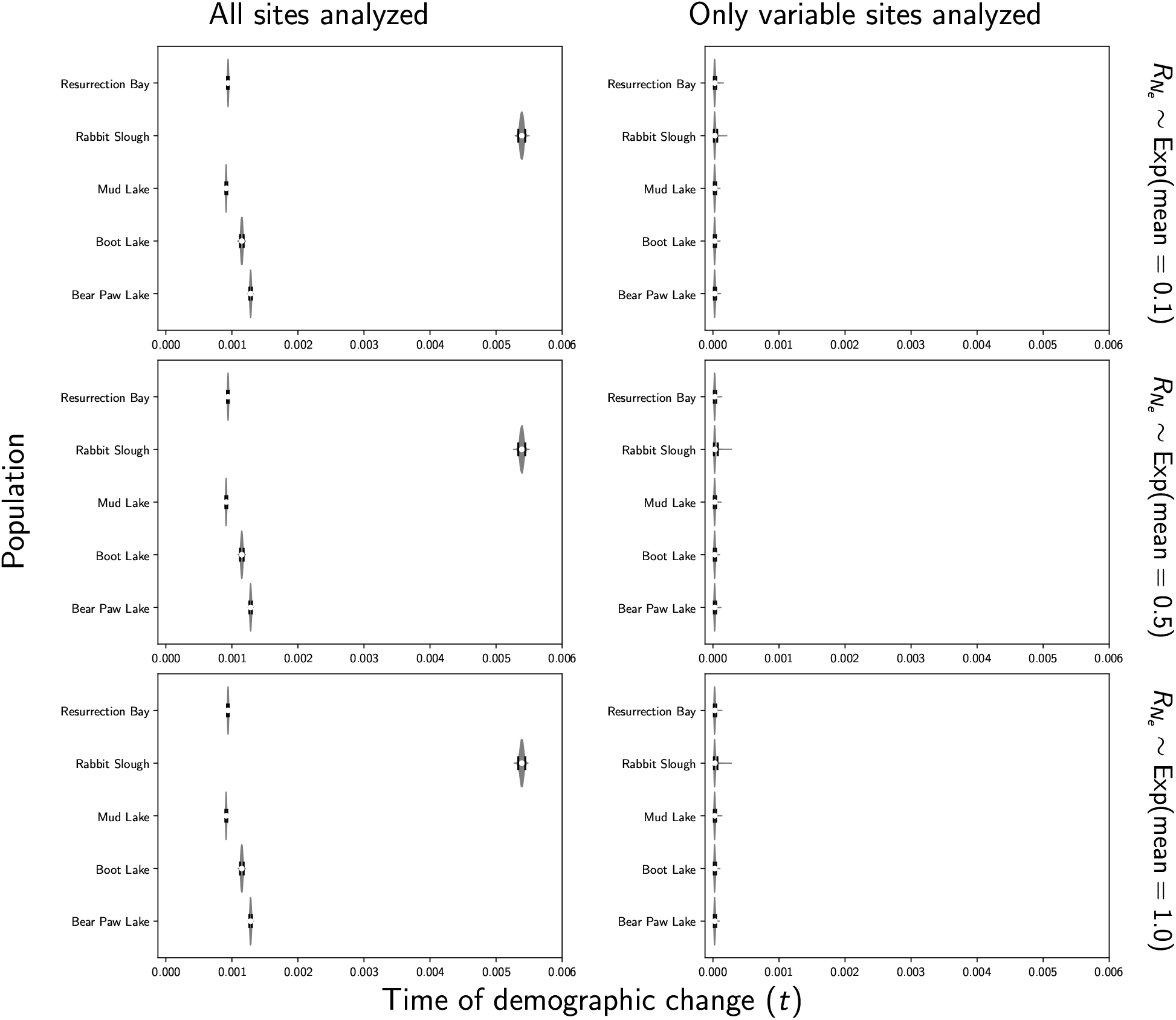
Estimates of the time of a change in population size across five stickleback populations when all of the sites (left column) or only variable sites (right column) of the RADseq alignments are analyzed. Each row shows results under a different prior on the relative effective size of the ancestral population. We generated the plots using matplotlib Version 2.0.0 (Hunter, 2007).

**Figure S24.**
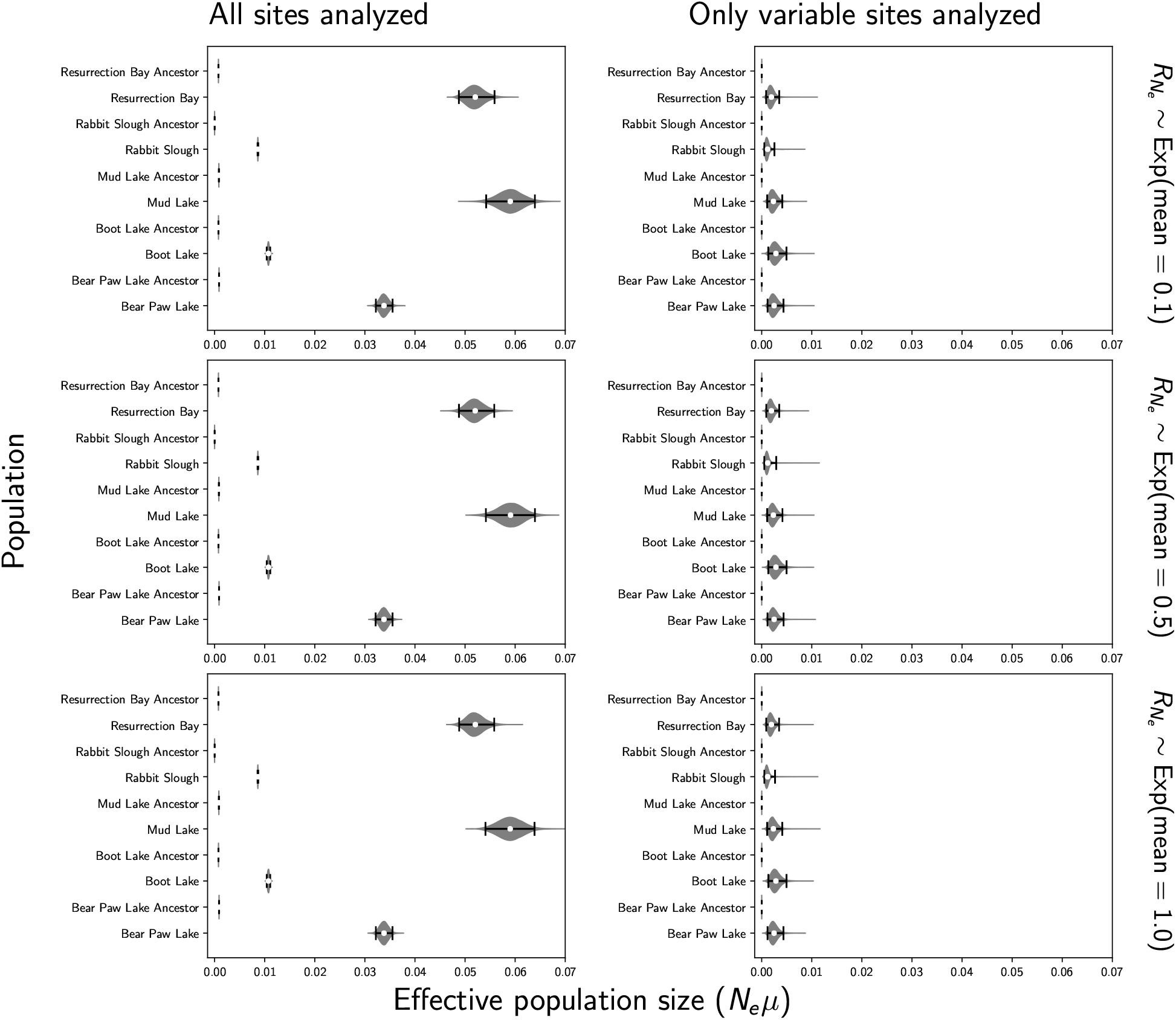
Estimates of the effective population size before (“ancestor”) and after a demographic change across five stickleback populations when all of the sites (left column) or only variable sites (right column) of the RADseq alignments are analyzed. Each row shows results under a different prior on the relative effective size of the ancestral population. We generated the plots using matplotlib Version 2.0.0 (Hunter, 2007).

